# Room-temperature crystallography reveals altered binding of small-molecule fragments to PTP1B

**DOI:** 10.1101/2022.11.02.514751

**Authors:** Tamar (Skaist) Mehlman, Justin T. Biel, Syeda Maryam Azeem, Elliot R. Nelson, Sakib Hossain, Louise E. Dunnett, Neil G. Paterson, Alice Douangamath, Romain Talon, Danny Axford, Helen Orins, Frank von Delft, Daniel A. Keedy

## Abstract

Much of our current understanding of how small-molecule ligands interact with proteins stems from X-ray crystal structures determined at cryogenic (cryo) temperature. For proteins alone, room-temperature (RT) crystallography can reveal previously hidden, biologically relevant alternate conformations. However, less is understood about how RT crystallography may impact the conformational landscapes of protein-ligand complexes. Previously we showed that small-molecule fragments cluster in putative allosteric sites using a cryo crystallographic screen of the therapeutic target PTP1B (Keedy*, Hill*, 2018). Here we have performed two RT crystallographic screens of PTP1B using many of the same fragments, representing the largest RT crystallographic screens of a diverse library of ligands to date, and enabling a direct interrogation of the effect of data collection temperature on protein-ligand interactions. We show that at RT, fewer ligands bind, and often more weakly -- but with a variety of temperature-dependent differences, including unique binding poses, changes in solvation, new binding sites, and distinct protein allosteric conformational responses. Overall, this work suggests that the vast body of existing cryogenic-temperature protein-ligand structures may provide an incomplete picture, and highlights the potential of RT crystallography to help complete this picture by revealing distinct conformational modes of protein-ligand systems. Our results may inspire future use of RT crystallography to interrogate the roles of protein-ligand conformational ensembles in biological function.

## Introduction

Of the ~150,000 protein crystal structures in the public Protein Data Bank (PDB) (Berman et al., 2000), ~122,000 (~81%) have a non-polymer ligand modeled, and many thousands more reside in private pharmaceutical company databases. However, of the trove of public protein-ligand crystal structures, the vast majority (~94%) with temperature annotations were determined at cryogenic (cryo) temperature (≤ 200 K), typically after the protein crystals were flash-cooled in liquid nitrogen. By contrast, only a small minority (~6%) were determined at elevated temperatures (> 200 K; of these, mostly > 277 K or 0°C). This statistic is unnerving in light of the fact that, for proteins, cryo crystallography distorts protein conformational ensembles (Keedy et al., 2014), whereas room-temperature (RT) crystallography reveals distinct protein conformational heterogeneity, including alternate conformations of side chains and backbone segments, that better aligns with solution data and is in some cases more relevant to biological function (Fenwick et al., 2014; Fraser et al., 2009, 2011; Keedy et al., 2015).

In contrast to proteins, relatively little is known about how crystallographic temperature affects protein-ligand interactions. Past studies that focused on individual compounds or small sets of related/congeneric compounds have offered tantalizing hints, with RT resulting in shifted binding poses (Bradford et al., 2021; Gildea et al., 2021; Maeki et al., 2020; Milano et al., 2022), binding at a different site (Fischer et al., 2015), and even a change of crystal symmetry (Gildea et al., 2021). In general, RT crystallography of protein-ligand complexes is increasingly accessible thanks to advances in methodology (Fischer, 2021) including serial crystallography (Milano et al., 2022), even with as few as 1000 images, at synchrotrons (Weinert et al., 2017) or X-ray free electron lasers (XFELs) (Moreno-Chicano et al., 2019). Other RT approaches are also emerging, including in situ crystallography (Lieske et al., 2019; Sanchez-Weatherby et al., 2019), in some cases using crystallization plates pre-coated with dry compounds (Gelin et al., 2015; Teplitsky et al., 2015), as well as microfluidics (Maeki et al., 2020; Sui et al., 2021).

Despite this promising foundation, a central question remains: how frequently, and in what ways, does temperature affect protein-ligand structural interactions? To our knowledge, this question has not yet been addressed using a sufficiently large library of chemically diverse ligands. This gap is a significant obstacle toward a thorough understanding of how ligand and protein conformational heterogeneity interplay with one another to control biologically important phenomena such as enzyme catalysis and allosteric regulation. It also limits the potential of structure-based drug design (SBDD), given that cryo temperature is reported to degrade the utility of crystal structures for computational docking and binding free energy calculations (Bradford et al., 2021).

An emerging high-throughput approach to identifying protein-ligand hits is crystallographic small-molecule fragment screening, in which hundreds to thousands of “fragments’’ of drug-like small molecules are subjected to high-throughput crystal soaking and structure determination with a protein of interest. For example, a recent crystallographic fragment screen of the SARS-CoV-2 coronavirus’s main protease (M^pro^) (Douangamath et al., 2020) provided dozens of starting points for crowd-sourcing the design of potent new small-molecule inhibitor candidates (The COVID Moonshot Consortium et al., 2022), complementing another crystallographic screen of M^pro^ using repurposed drug molecules (Günther et al., 2021).

Previously, several authors of the current study performed a crystallographic fragment screen of the archetypal protein tyrosine phosphatase, PTP1B (also known as PTPN1) (Keedy et al. 2018), a highly validated therapeutic target for diabetes (Elchebly et al., 1999), cancer (Krishnan et al., 2014), and neurological disorders (Krishnan et al., 2015) that has also been deemed “undruggable” (Mullard, 2018; Zhang, 2017). That fragment screen produced X-ray datasets for 1,627 unique fragments, of which 110 were clearly resolved in electron density maps at 12 fragment-binding sites scattered across the surface of PTP1B. Of the top three fragment-binding “hotspots”, one was previously validated with a noncovalent allosteric small-molecule inhibitor (Wiesmann et al., 2004), and another was validated with a new covalent allosteric inhibitor inspired by the fragment hits (Keedy et al., 2018), thus highlighting fragment screening as a valuable tool for discovering allosteric footholds in proteins (Krojer et al., 2020). Importantly, however, all previously reported crystallographic fragment screens, including those mentioned above, were conducted at cryogenic (cryo) temperature.

To elucidate the role of temperature in dictating protein-ligand interactions, here we have explored the use of large-scale crystallographic small-molecule fragment screening at room temperature. Specifically, we have performed RT crystallographic fragment screens of PTP1B with many of the same fragments used for the previous cryo screen (Keedy et al., 2018), thereby allowing direct inferences regarding the effects of temperature. Our work uses 143 unique, chemically diverse ligands which, to our knowledge, is several-fold (4-5x) more than any previous RT crystallography study. Moreover, we have used two complementary strategies for RT data collection. In the first, each RT dataset derives from a single crystal after manual harvesting and mounting on a nylon loop, as with most traditional crystallography (including the previous cryo fragment screen of PTP1B). In the second, each RT dataset derives from several crystals subjected to diffraction in situ in the original mother liquor and crystallization plate. The two screens were performed with different diffraction data collection approaches, at different times, and with distinct but partially overlapping sets of fragments -- so together they ensure that our overall conclusions are robust.

In both RT fragment screens, we observe that fewer fragments bind, and on average more weakly (with lower occupancy). However, many fragments that bind at RT do so with a variety of temperature-dependent differences, including unique binding poses, changes in solvation, totally new binding sites, and even distinct protein allosteric conformational responses (Choy et al., 2017; Cui et al., 2017; Hjortness et al., 2018; Hongdusit et al., 2020; Keedy et al., 2018; Torgeson et al., 2022) to ligand binding. Serendipitously, we also identify a fragment that binds covalently to a key lysine in the allosteric 197 site (Keedy et al., 2018), providing an intriguing new foothold for further allosteric inhibitor development.

Overall, this work provides new insights into ligandability and allostery in the important therapeutic target enzyme PTP1B. More broadly, it highlights the limitations of relying solely on cryo crystallography and the relative advantages of room-temperature crystallography for elucidating interactions between ligands and proteins, with implications for a wide range of applications including structure-based drug design.

## Results

### Two crystallographic fragment screens at room temperature

This work centers on two room-temperature crystallographic screens of PTP1B: single-crystal (hereafter abbreviated as “1-xtal”) and in-situ. For each RT screen, we used small-molecule fragments that were used previously for the cryogenic-temperature crystallographic screen (Keedy et al., 2018) (see Methods). Fragments were chosen from two categories: (1) *cryo-hits* which bound to the protein in the previous cryo screen, and (2) *cryo-non-hits* which were soaked into crystals but did not bind in the previous cryo screen. The 1-xtal RT screen used 59 cryo-hits and 51 cryo-non-hits, whereas the in-situ RT screen used 48 cryo-hits and 32 cryo-non-hits. The fragment sets for the two screens were partially overlapping and complementary, with 23 fragments in common, of which 20 were cryo-hits and 3 were cryo-non-hits.

The fragment-soaked and control dataset for both screens are totaled and categorized in **Table 1**. Unless noted otherwise, the unique fragment datasets plus DMSO datasets for each screen were used for all subsequent analyses. As the two screens had 23 fragments in common, there were a total of 86 + 80 - 23 = 143 unique fragments overall across both RT screens.

**Table 1:** X-ray datasets collected for both room-temperature crystallographic screens. The total datasets tally for the in-situ screen derives from a larger number of partial datasets or “wedges” that were merged (see Methods). The cryo-hit and cryo-non-hit categories are defined in Methods. Datasets from crystals soaked with DMSO only are included in the total datasets tally, but not in the unique fragments tally.

|  | 1-xtal | In-situ |
| --- | --- | --- |
| Total raw X-ray datasets collected | 269 | 111 |
| Processed datasets with unique fragments | 86 | 80 |
| Cryo-hit | 38 | 48 |
| Cryo-non-hit | 48 | 32 |
| DMSO (negative control) | 7 | 20 |

The data were high-resolution for both RT screens (**Fig. 1**): the average resolution was 2.30 Å for 1-xtal and 1.99 Å for in-situ, as compared to 2.10 Å for the previous cryogenic screen (Keedy et al., 2018). The slightly lower resolution of the 1-xtal data may be due to some degree of radiation damage, which was largely avoided by the in-situ strategy (see Methods). As outlined below, the results of the two screens are broadly very similar, and indeed identical for several fragments used in both screens (**Fig. S7**), suggesting that radiation damage with the 1-xtal data was not a major factor in dictating our overall results. Additionally, a visual inspection of all the 1-xtal RT hits featured in this paper did not show any signs of local radiation damage (see Methods).

**Figure 1:**
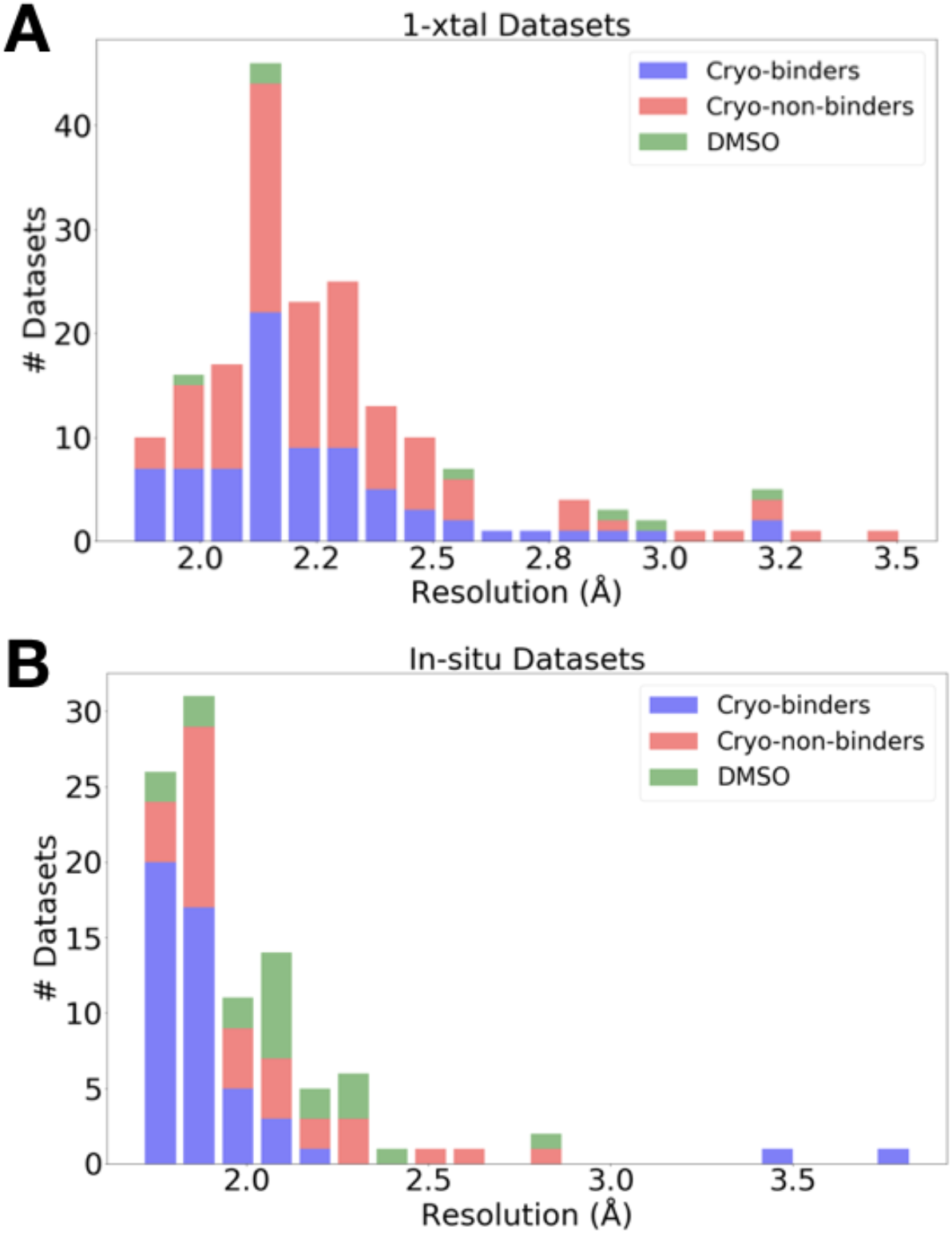
Resolution distributions from room-temperature crystallographic screens. Histogram of X-ray resolutions of datasets soaked with DMSO (green), cryo-hits compounds (blue), or cryo-non-hits (red), collected at RT via **(A)** 1-xtal or **(B)** in-situ data collection techniques.

### Identifying fragment binding hits

Using these high-resolution datasets, we identified low-occupancy protein-fragment binding events using the PanDDA algorithm (Pearce, Krojer, Bradley, et al., 2017) and manual inspection and modeling (see Methods). For the fragments that bound to PTP1B at cryo (cryo-hits) (Keedy et al., 2018), we then examined how many bound to PTP1B at RT. The initial hit rates from the event maps automatically generated by PanDDA were surprisingly low: 12/38 (32%) for 1-xtal and 7/48 (15%) for in-situ. Additionally, for cryo-non-hits, PanDDA revealed only 2 binding events; both were for the same fragment in the same dataset (*vide infra*).

To identify hits that may have been missed by the automated PanDDA event identification algorithm, we manually generated RT event maps with the cryo value for 1-BDC, a quantity within PanDDA that is directly related to ligand binding occupancy (Pearce, Krojer, Bradley, et al., 2017) (see Methods). With this approach, we found 5 new binding events: 3 for the 1-xtal datasets and 2 for in-situ datasets. This brought the new totals to 15/38 (39%) for 1-xtal and 9/48 (19%) for in-situ, still fairly low hit rates.

This observation prompted us to reexamine how the many partial datasets or “wedges’’ obtained from in-situ crystallography are assembled into complete datasets for use in subsequent steps including map calculation and PanDDA modeling (see Methods). Recently, a new software called cluster4x was unveiled for pre-clustering X-ray datasets in the space of differences in structure factor amplitudes and/or Cα positions (Ginn, 2020). When applied to our past cryogenic PTP1B screen (Keedy et al., 2018), cluster4x identified previously unrecognized binding events (Ginn, 2020). The RT datasets are more isomorphous than the past cryo datasets (**Fig. S1**). Nevertheless, to enhance isomorphism for our in-situ screen, here we used cluster4x to pre-cluster in-situ wedges (**Fig. S2**) before merging within 3 main clusters (**Table S1**) that are partially overlapping but qualitatively distinct from each other. We then assembled sets of similar wedges into complete datasets (see Methods) for input to PanDDA. This pre-clustering protocol resulted in 5 additional hits that were not previously observed with the all-wedges datasets, bringing the total RT hit rate for cryo-hits up from 9/48 (19%) to 14/48 (29%) for in-situ.

Given the final cryo-hit reproduction rates of 39% and 29% for the RT screens, we investigated whether temperature affected the binding occupancy, or percent of unit cells in the crystal with a fragment bound. As an accessible proxy for occupancy, we examined PanDDA 1-BDC values. Many fragments have lower occupancy at RT than at cryo (**Fig. 2**). This trend holds for cryo-hits that bind to the cryo site at RT either with the same pose (blue points in **Fig. 2**) or with a new pose (orange points in **Fig. 2**) (see **Table 2** below). The lower occupancies at RT suggest that our cryo-hit reproduction rate at RT (mentioned earlier; also indicated as gray points in **Fig. 2**) is likely due to changes in fragment binding as a function of temperature rather than e.g. experimental errors.

**Figure 2:**
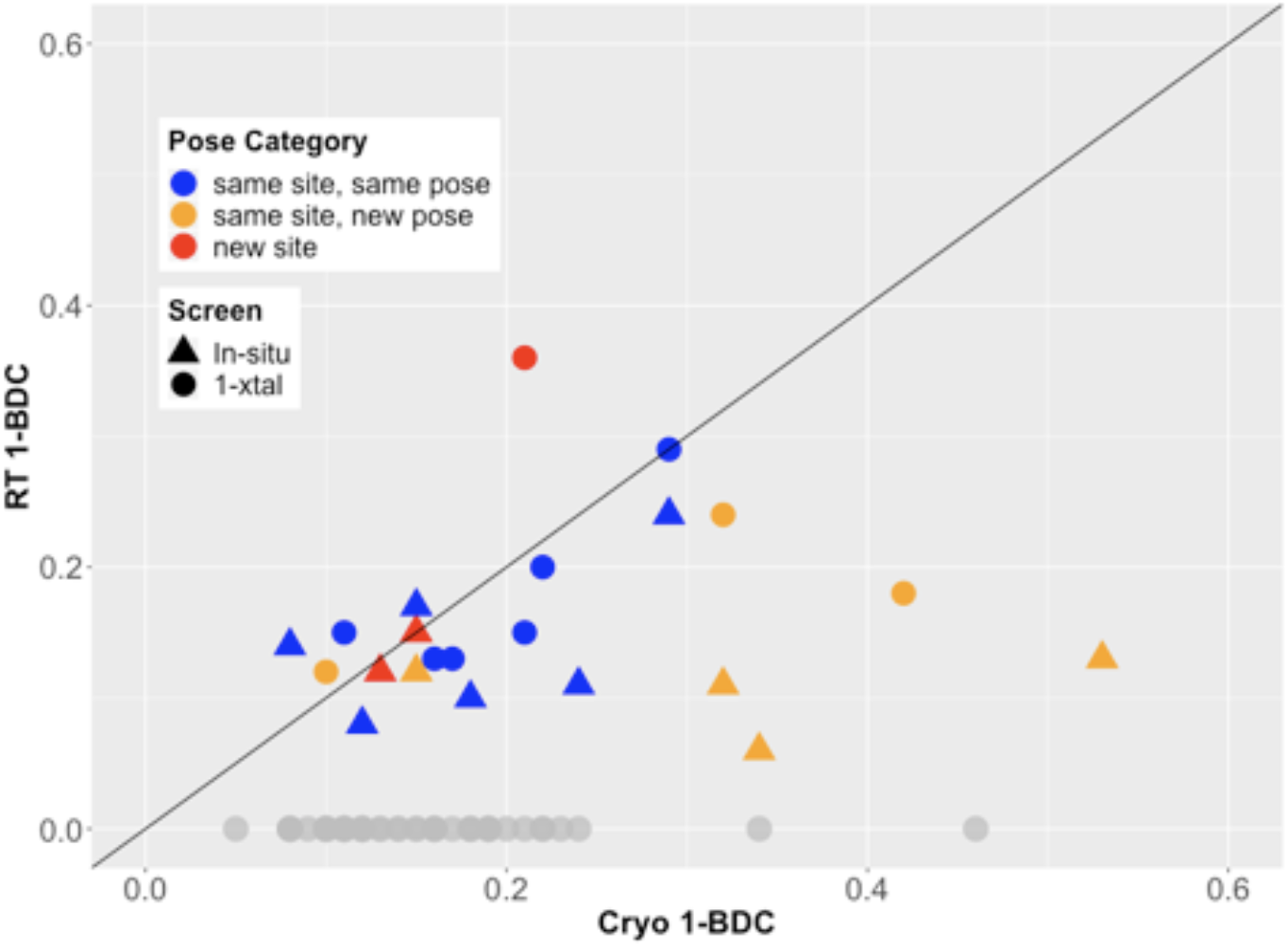
Fragment binding occupancies are different and often lower at room temperature. 1-BDC (a proxy for occupancy) is plotted for each binding event observed in either of two RT screens vs. in the previous cryo screen. For 2 datasets, 2 binding events for the same fragment in the same structure are included as separate points. See **Table 2** for definitions of pose categories. Those that did not show binding at RT are in gray along the x-axis. In some additional cases, RT event maps were calculated using the cryo 1-BDC to identify bound ligands at RT; these cases would sit artificially on the diagonal, and are not shown here.

**Table 2:** Characteristics of fragment hits for room-temperature screens. For each of the two RT screens, 1-xtal and in-situ, all modelable fragment binding events are categorized on the basis of the binding pose relative to the cryo-temperature binding pose. The tallies are on the basis of individual fragment binding events, not crystal structures containing the fragment, because in some cases a given fragment binds at multiple sites, e.g. both the cryo-hit site and a new site. Thus, the “new site” category includes both cryo-hits that bind at a new site and cryo-non-hits that now bind at RT. The categories are not all mutually exclusive: for example, some fragments bind with the same site and same pose, but induce significant protein conformational change. * The “same site, new pose” category includes cases in which the fragment pose is the same but the protein conformation is notably altered at RT vs. cryo. See **Table S2** for more information on which individual datasets fit into which categories.

| RT pose category | 1-xtal | In-situ | Total |
| --- | --- | --- | --- |
| same site, same pose | 9 | 7 | 16 |
| same site, new pose* | 4 | 5 | 9 |
| new site | 3 | 2 | 5 |
| protein change | 2 | 3 | 5 |
| not hit | 71 | 66 | 137 |

### Distribution of fragment hits at room temperature

Between our two RT screens, we have 15 + 14 = 29 new room-temperature events with small-molecule fragments. These fragments fall into several categories based on the binding site, binding pose, and conformational response by the binding site (**Table 2**, **Table S2**).

The RT fragment hits are bound at sites distributed throughout PTP1B, including at least one at the active site and at all three allosteric sites previously highlighted by the cryo screen (Keedy et al., 2018): the 197 site, BB site, and L16 site (**Fig. 3**, **Fig. S3**). Many of these sites have RT hits in both the 1-xtal and in-situ screens (**Fig. S3**), confirming the success of both screens. Notably, in all of these four key sites, one or more fragments bind differently from cryo -- either binding with a new pose at RT, or binding to this site only at RT and not cryo (**Fig. 3**).

**Figure 3:**
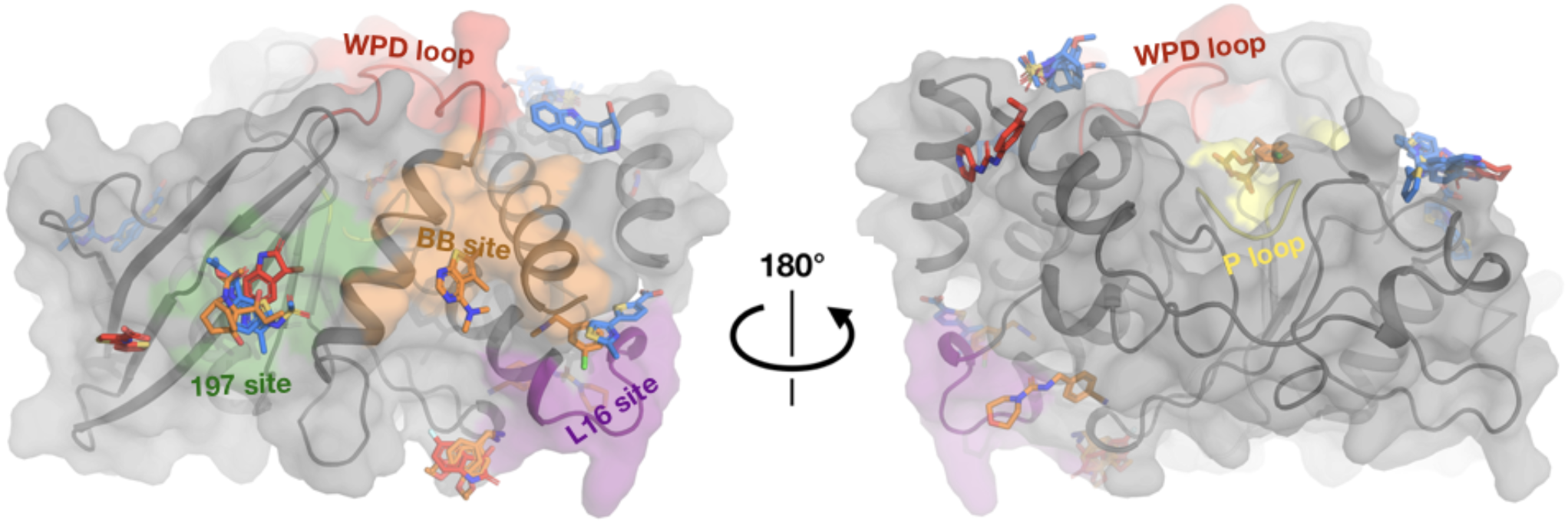
Fragments have a similar distribution across protein sites but different binding modes at room temperature. Overview of fragments bound across PTP1B at RT, colored by RT pose compared to cryo pose: same site, same pose (blue); same site, new pose (orange); new site (red). See **Table 2** for more details on the definitions of these classifications. Also highlighted are the active-site WPD loop (red), P loop (yellow) 197 allosteric site (green), BB allosteric site (orange), and L16 allosteric site (purple) (Keedy et al., 2018). The protein is shown in its open conformation with the WPD loop and L16 in the open state. The α7 helix is not shown since it is disordered when the protein is in the open state, which is favored at higher temperatures (Keedy et al., 2018). α7 does become ordered in one RT fragment-bound structure, but is not shown here.

### Similar binding for many cryo-hits at room temperature

We next turned our attention to the precise binding poses of cryo-hit fragments at RT. Of the cryo-hits that also bind at RT, most do so with a similar pose (**Fig. 4**, **Table 2**): 9 cases for 1-xtal (**Fig. S4**) and 7 for in-situ (**Fig. S5**). Many of these are concentrated in two sites on the non-allosteric front side of the protein (**Fig. S3**) that were also highly populated in the cryo screen. Additionally, some fragments are double represented due to the overlap between the two screens. Notably, of the 3 fragments with binding events in both the 1-xtal and in-situ screens, all 3 bind similarly in both RT screens (**Fig. S7)**, suggesting the RT results are reproducible and reliable.

**Figure 4:**
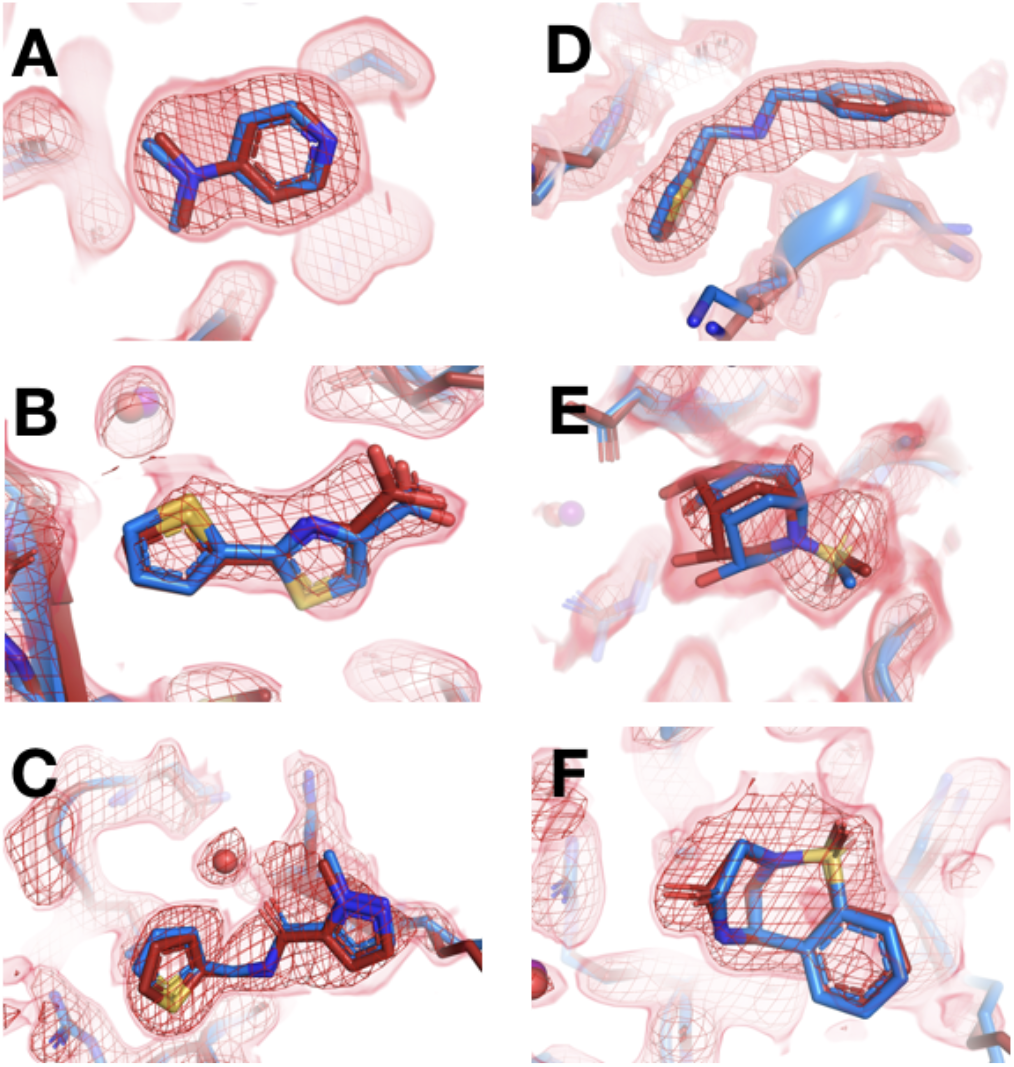
Fragments that bind similarly at room vs. cryo temperatures. For each dataset, the RT PanDDA event map is in red (contour levels below), the RT model is in red (waters in red), and the corresponding cryo model is in blue (waters in purple). Datasets are named as follows: x#### = RT in-situ, z#### = RT 1-xtal, y#### = cryo. **(A-C)** in-situ. **(D-F)** 1-xtal. **(A)** RT: x0224 (2.0 σ), cryo: y0118 **(B)** RT: x0285 (1.5 σ), cryo: y0772 **(C)** RT: x0262 (1.5 σ), cryo: y1656 **(D)** RT: z0007 (2.0 σ), cryo: y1710 **(E)** RT: z0015 (1.8 σ), cryo: y1554 **(F)** RT: z0025 (1.5 σ), cryo: y1294

This figure contains selected examples of fragments that bind similarly at RT vs. cryo; for all examples, see **Fig. S4** for 1-xtal and **Fig. S5** for in-situ. For examples with no RT density for the cryo ligand using the cryo 1-BDC, see **Fig. S6**.

Although all of the aforementioned fragments themselves bind with the same pose at RT vs. cryo, in some cases water molecules around them differ with temperature. In a few examples, clear event map density is present for a water at RT but not at cryo **(Fig. 4C**, **Fig. S8A-B**) or vice versa (**Fig. S8C)**, even when varying event map contour levels. Therefore, even when ligand binding is similar, the solvation layer around the ligand can change at cryo vs. RT.

### New binding poses at room temperature

Some cryo-hit fragments bind in the same site at RT, but with a quite different pose. In one striking example, the fragment binds with the central ring in the same position at RT vs. cryo, but with substantially different positions for the two substituent groups (**Fig. 5**). The in-plane chlorine and out-of-plane methylamine group are clearly defined in the respective event maps: the RT density is incompatible with the cryo model, and vice versa. Notably, this fragment binds in the allosteric L16 site (**Fig. 3**), which was first reported alongside the original cryo fragment screen for PTP1B and highlighted as a promising target for small-molecule allosteric inhibitor development (Keedy et al., 2018). Thus, RT crystallography can reveal novel ligand poses in allosteric sites, with possible relevance for elucidating allosteric mechanisms and informing allosteric inhibitor design.

**Figure 5:**
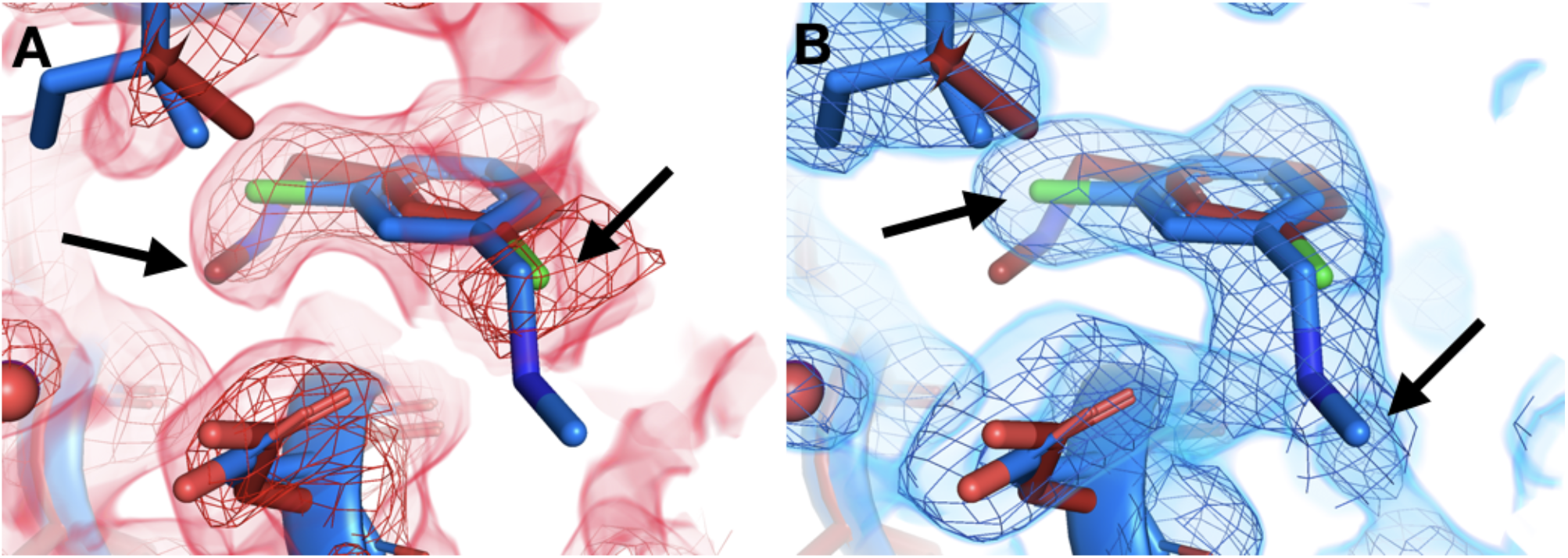
Temperature-dependent ligand conformational heterogeneity. **(A)** In the RT dataset (x0227), the RT event map contoured at 2 σ (red) matches the RT model (red) rather than the cryo model (blue) for both substituent groups of the ring. **(B)** In the cryo dataset (y0071), the cryo event map contoured at 1.2 σ (blue) matches the cryo model rather than the RT model.

Another example features alternate ligand conformations that coexist in the same site, but only at one temperature. The RT event map suggests a pose with the carbonyl pointed one direction, toward Arg238 (**Fig. 6A**). However, at cryo, this fragment was previously modeled with the carbonyl rotated by a 180° flip, enabling a water-bridged H-bond (**Fig. 6B**). The RT event map has weak evidence at best for the flipped cryo conformation (**Fig. 6A**). By contrast, the cryo event map has significant evidence for both conformations (**Fig. 6B**). This observation is akin to other examples in which a ligand exhibits conformational heterogeneity in a single X-ray dataset (van Zundert et al., 2018). Here, however, the ligand conformational heterogeneity is temperature-dependent, enabling cross-pollination of conformations across temperatures to improve modeling (Bradford et al., 2021; Keedy, 2019).

**Figure 6:**
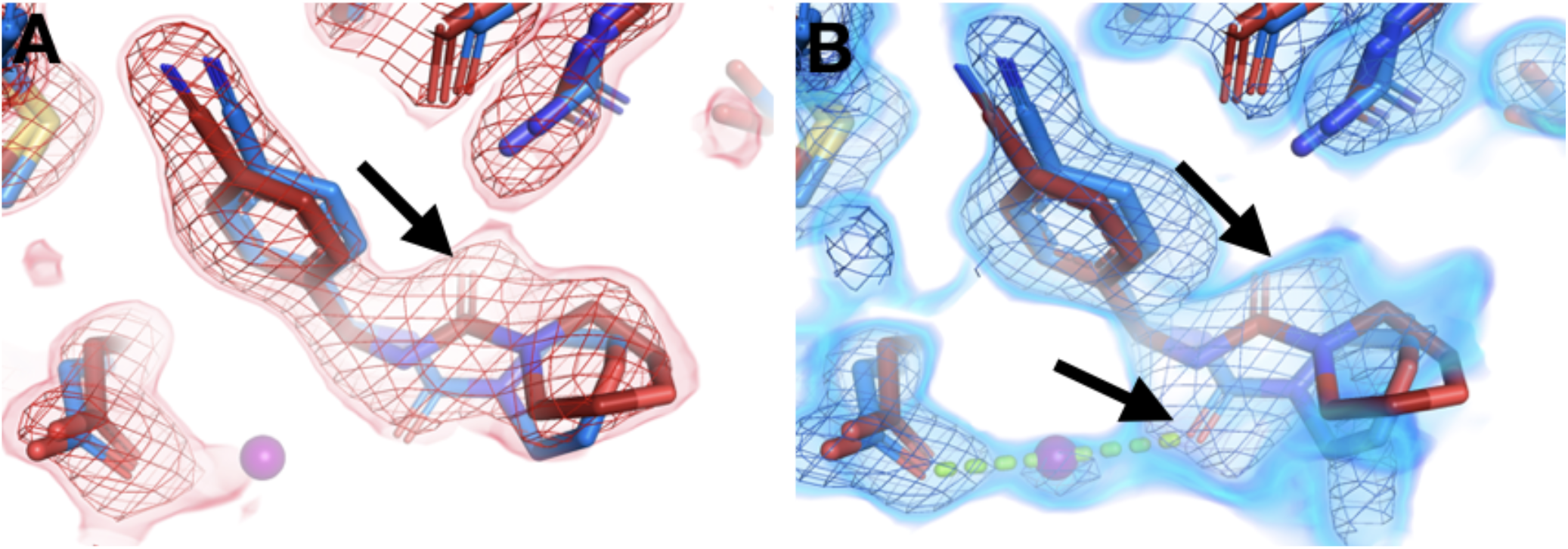
Temperature modulates fragment pose and solvation within the same site. **(A)** In the RT dataset (x0260), the RT event map contoured at 1.6 σ (red) matches the RT model (red), but shows little evidence for the cryo model (y0180, blue). **(B)** In the corresponding cryo dataset, the cryo event map contoured at 1.6 σ (blue) matches both the cryo model (blue) and the RT model. Notably, only at cryo does the event map include density for a water molecule (purple ball) next to the fragment carbonyl group and well-positioned for a hydrogen bond (pale green dashed line) with the cryo fragment pose.

A pair of other examples also feature fragments with distinct poses that are differentially stabilized at RT vs. cryo. In each of these two related examples, the RT event density is clear that the fragment binds with its longer substituent well-ordered and pointed underneath the active-site WPD loop, which closes over the fragment (**Fig. 7A,B**, left panels). At cryo, the loop still closes over the fragment, and the core of the fragment is in a similar location. However, the cryo event density is inconsistent with the RT pose -- instead, the longer substituent seems to protrude toward solution (**Fig. 7A,B**, right panels). For one of these fragments, a new ordered water molecule at cryo displaces the RT ligand pose (**Fig. 7B**).

**Figure 7:**
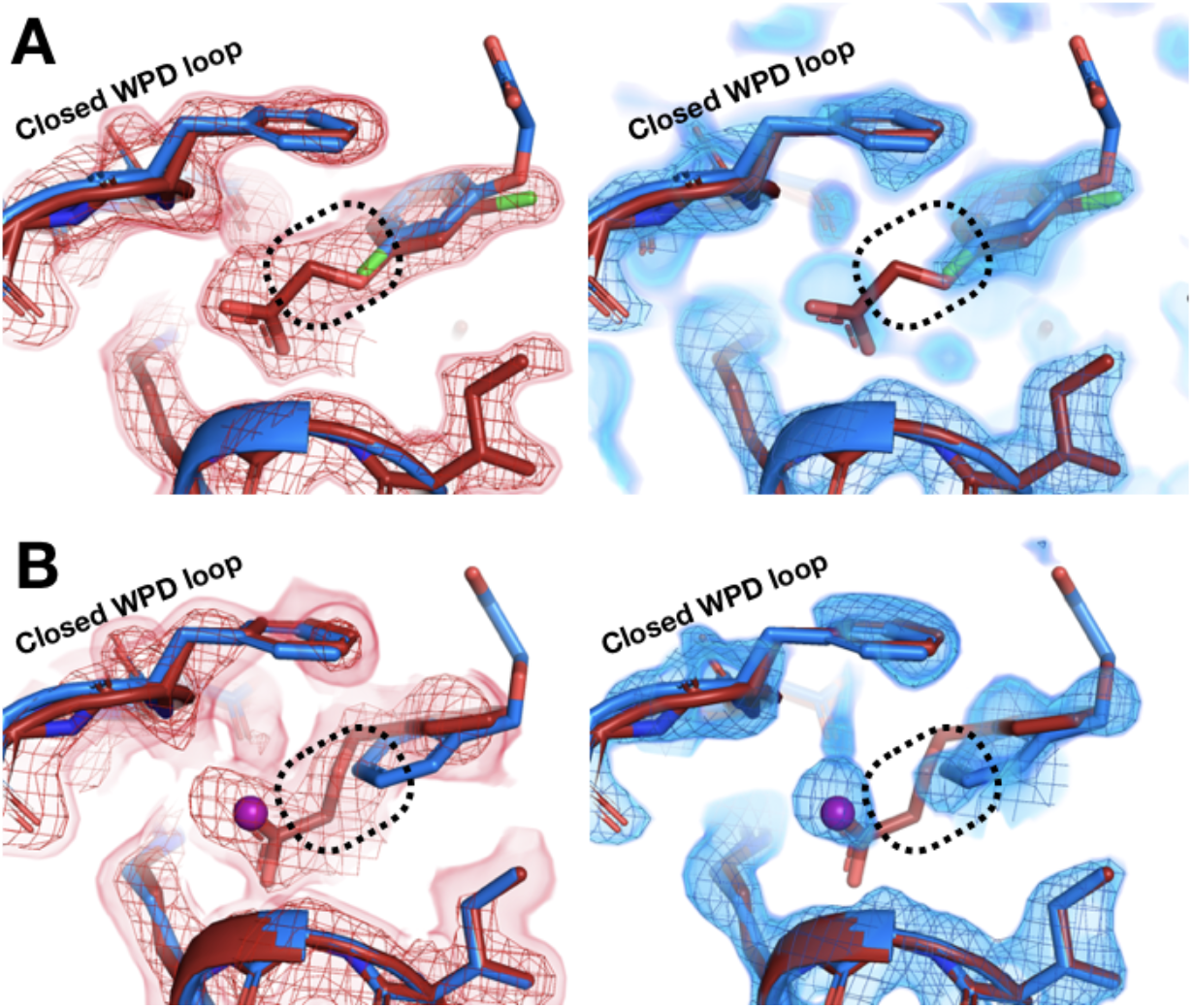
Room-temperature fragment pose is flipped compared to the cryo pose. **(A)** *Left:* RT density (red) 1.5 σ, z0055 (red); y0884 (blue). *Right:* cryo density (blue) 1 σ, z0055 (red); y0884 (blue). **(B)** *Left:* RT density (red) 2 σ, x0256 (red); y0650 (blue). *Right:* cryo density (blue) 0.8 σ, x0256 (red); y0650 (blue) (not previously deposited to the PDB). Density is linked at RT (dashed box), consistent with the fragment pose, but is cut off at cryo, even at lower contour. There is little to no density for the open state of the WPD loop (not shown).

### New binding sites at room temperature

Beyond just differences within the same binding site, temperature can also modulate ligand binding more dramatically, even altering what protein site the ligand binds to. In a first example, the fragment binds to the allosteric BB site (Keedy et al., 2018; Wiesmann et al., 2004) at cryo (**Fig. 8A,C**), but there is no event density at RT. Instead, there is strong fragment binding event density at a different site nearly 40 Å away (**Fig. 8A,B**). The RT event density supports subtle protein shifts in the new binding site to accommodate the new fragment binding event (**Fig. 8B**). By contrast, in the cryo binding site, the RT protein conformation would clash with the cryo pose, disallowing binding at RT.

**Figure 8:**
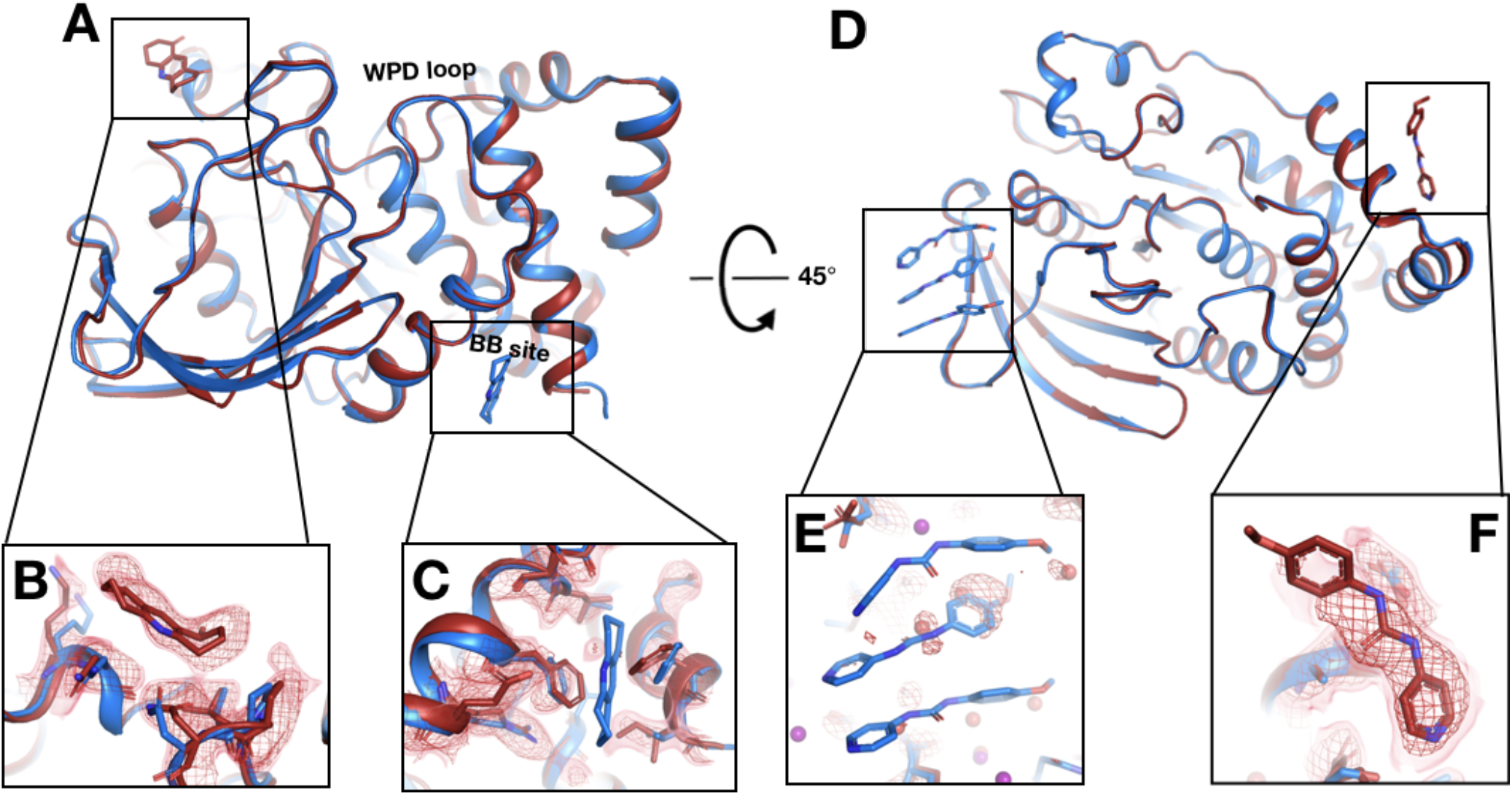
Fragments bind at new sites only at room temperature. **(A-C)** First example. **(A)** The two sites are ~38 Å away from one another. **(B)** In the RT dataset (z0042), the RT event map, calculated with 1-BDC of 0.36 and contoured at 1.5 σ (red), supports a bound fragment in the RT model (red) at a new site while the cryo model (y1525) (blue) has no bound fragment. **(C)** By contrast, the RT event map (same contour) does not show any density for the cryo model (blue) from the previous cryo dataset (y1525). **(D-F**) Second example. **(D)** The two sites are ~46 Å away from one another. **(E)** The RT event map contoured at 1.75 σ (red) (same contour) does not support the cryo model (blue) from the previous cryo dataset (y0572). **(F)** By contrast, at a new site the RT event map (same contour) supports a bound fragment in the RT model (x0225) (red). The cryo model has no bound fragment.

In additional examples, elevated temperature dissipates what seem to be cryo binding artifacts. In the first such example, at cryo the fragment binds with an artifactual stacking arrangement involving three copies of the fragment (**Fig. 8D,E**), but at RT there is no event density (automated or custom) for this stacking. This result suggests that temperature can modulate protein-ligand energy landscapes significantly, in this case by disfavoring enthalpically favorable stacking at higher temperature. Moreover, at RT, new event density for a single copy of this fragment appears at a distal site (**Fig. 8F**) that is over 45 Å away from the cryo site (**Fig. 8D**). Cryo event density at the new site was too weak to justify modeling a bound fragment (Keedy et al., 2018). Thus, the cryo binding site is unique to cryo and the RT binding site is unique to RT. In fact, this is the only case in which a fragment binds at RT to a new site that was not previously thought to bind any fragments at cryo (although later computational reanalysis did discover one previously undetected adjacent cryo-hit in this area (Ginn, 2020)). A Tris buffer molecule also fortuitously binds in the same location in another published structure (PDB ID 4y14), although it is held in place by a distinct crystal contact due to that structure’s space group.

In a related but distinct case, a fragment previously bound at cryo with a seemingly similar artifactual stacking arrangement, this time involving two copies of the fragment (**Fig. S9A**). However, at RT the entire stack does not disappear -- instead, one copy remains bound (**Fig. S9B**). At cryo, this latter copy was slightly more ordered than the other, based on event map strength. Thus, elevated temperature is sufficient to displace the more weakly bound copy, but not the more tightly bound one.

In a final, somewhat more complicated example, a fragment previously bound at three distal sites at cryo. At RT the fragment binds to only one of the cryo sites, in a nearly identical pose. In the other cryo sites, RT binding events were not readily detected, either automatically by PanDDA or in RT event maps calculated at the cryo events’ 1-BDC values. More strikingly, at RT the fragment now binds to an additional new site (**Fig. S10A,B**) that is over 40 Å away from any of the three cryo sites (**Fig. S10C**). Although fragment binding was clear in cryo event maps at the three cryo sites, cryo density was unconvincing at the RT site; therefore, no binding event was detectable at this new site at cryo. Thus, as with the examples above (**Fig. 8**), this fragment binds uniquely to a new site at RT.

### New covalent binding events to lysines

In addition to the fragments that switch binding sites at RT as detailed above, one fragment binds only in our RT datasets -- and in an unexpected fashion. In RT event maps, we observe strong event density at/near both the allosteric 197 and L16 sites (Keedy et al., 2018). Surprisingly, at each site, the event density is contiguous with the side chains of a nearby lysine residue (**Fig. 9**), consistent with covalent binding by the isatin-based fragment. First, at the allosteric L16 site, the fragment binds covalently to Lys237 (part of the eponymous Loop 16) -- although it binds adjacent to the L16 pocket itself, nearer to the allosteric BB site (**Fig. 9C**). Second, at the allosteric 197 site site, the fragment binds covalently to Lys197 with a pose that is strikingly similar to that of a covalent allosteric inhibitor tethered to a K197C mutant (Keedy et al., 2018) (**Fig. 9A**).

**Figure 9:**
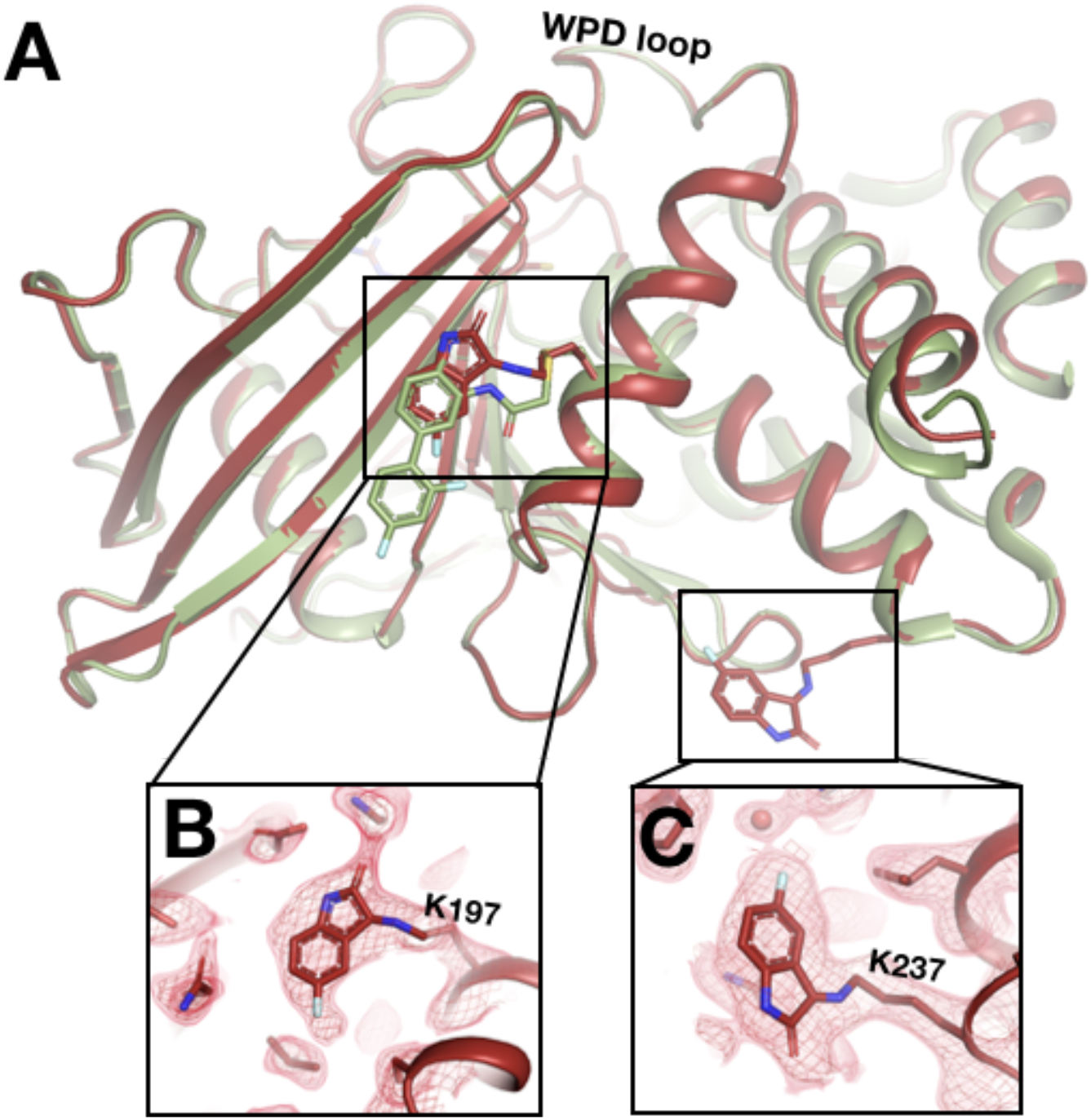
Unanticipated covalent adducts at previously reported allosteric sites only at room temperature. **(A)** RT structure with the fragment covalently bound to both K197 and K237 (z0048, red), aligned with cryo structure with a previously reported allosteric inhibitor covalently bound to K197C (6b95, green). **(B)** Fragment bound to K197 at the 197 allosteric site, with RT event density at 1.5 σ. **(C)** Fragment bound to K237 at the L16 allosteric site, with RT event density at 1.5 σ.

The distal active-site P loop and substrate-binding loop adopt new conformations that are similar to those observed when the catalytic Cys215 is oxidized (van Montfort et al., 2003), although it is unclear whether Cys215 is oxidized in our RT event map. These conformations were not observed with the K197C-tethered allosteric inhibitor (**Fig. S11**).

This fragment is a cryo-non-hit, meaning it demonstrably did not bind at cryo despite a high-resolution cryo dataset (1.89 Å, y1159). Indeed, it is the only cryo-non-hit to bind in either RT screen. This cryo-non-hit was chemically dissimilar to all previous cryo-hits: the most similar cryo-hit has a low Tanimoto score relative to this RT fragment (0.36, y1703) and does not bind near the RT sites. It is possible that the crystal for the cryo dataset was insufficiently soaked with this compound, or that the new RT binding events seen here are due to additional chemical changes to the compound in DMSO solvent over time that altered its reactivity toward lysines. As expected for fragments due to their weak binding affinities, this molecule does not inhibit PTP1B with an in vitro activity assay (Keedy et al., 2018) (data not shown). However, our observations here raise the hope that optimized versions of this compound, particularly driven by fragment linking of the K197C-targeted compound and this new fragment (**Fig. 9A**), could yield potent allosteric inhibitors for wildtype PTP1B, without need for mutation to a cysteine.

### Unique protein conformational responses at room temperature

Temperature does not only affect fragment binding to the protein -- it can also affect the protein’s conformational response to fragment binding. With both screens, we observe protein conformational responses that are preferentially localized to the key allosteric sites that were identified in our previous study as being inherently linked to the active site (Keedy et al., 2018).

The C-terminal end of the α6 helix forms part of the allosteric L16 site (Keedy et al., 2018). At cryo, fragments in this site that intercalate below the α6 helix push it further in the direction of α7, the BB site, and the rest of the allosteric network (Keedy et al., 2018). At RT, structures with two of these fragments (**Fig. S5G**, **Fig. 5**) show that they affect the position of α6 similarly at RT vs. cryo (**Movie S1**); perhaps surprisingly, this remains true despite one fragment exhibiting a 180° pose flip (**Fig. 5**).

However, in the nearby allosteric BB site (Wiesmann et al., 2004), the α6 helix is differentially ordered upon binding of a fragment at RT vs. cryo **(Fig. 10).** Although the fragment binds in the same pose at RT and cryo, an entire additional helical turn of α6 is ordered at RT. This example illustrates that temperature can modulate not only the positions of protein structural elements during ligand binding, but also their relative order vs. disorder.

**Figure 10:**
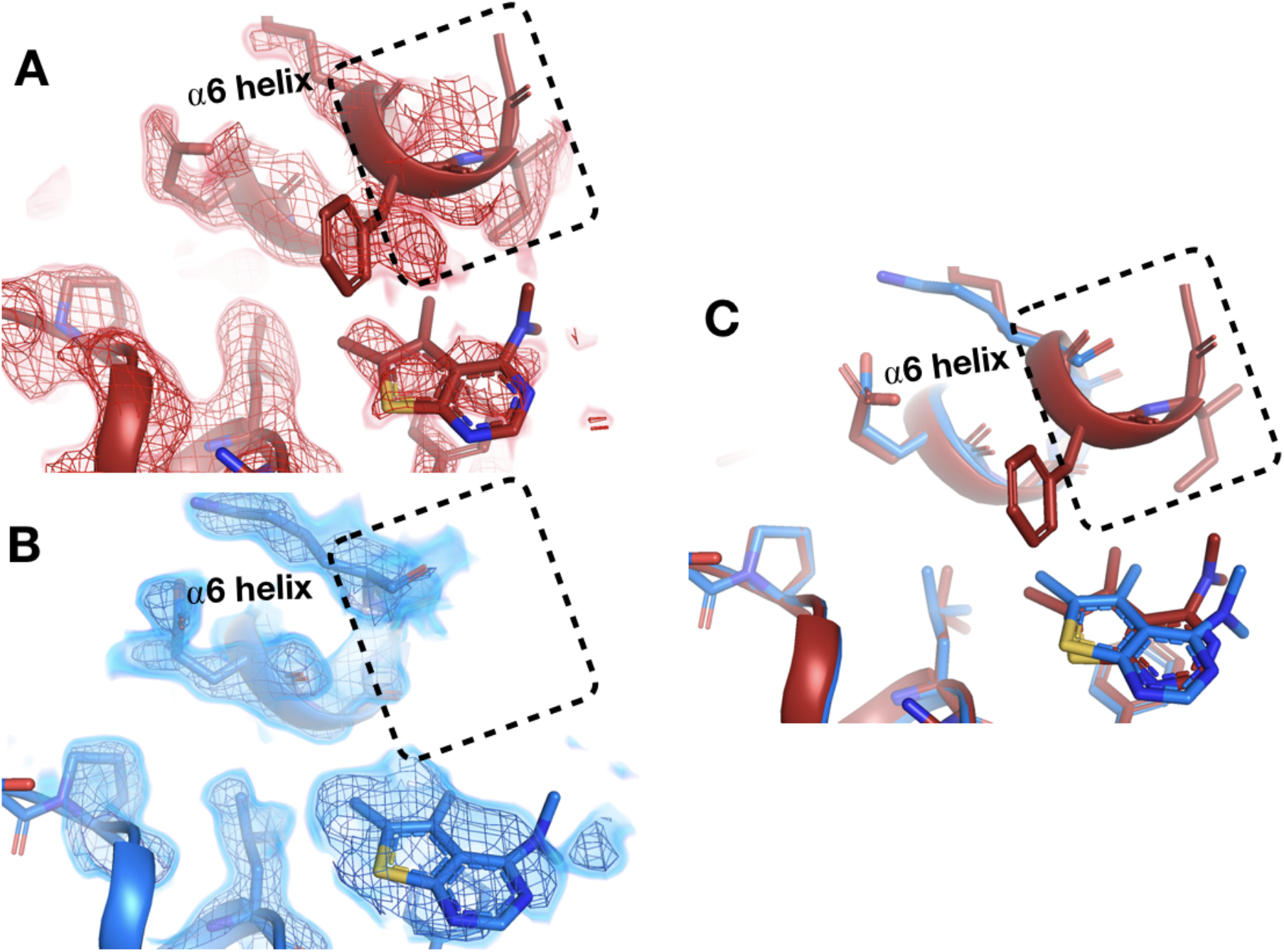
Temperature-dependent ordering of an α-helix augmenting a fragment binding site. **(A)** In the BB allosteric site, the RT density, x0222 (red); contoured at 1.25 σ, is consistent with an extended and more ordered α6 helix (dashed box). **(B)** In contrast, the cryo density, y0205 (blue); contoured at 1.75 σ, becomes disordered and therefore the α6 helix is not modeled as extended as in the RT model (dashed box). **(C)** Overlay of the two models showing the fragment pose is extremely similar whereas the RT helix is extended and more ordered (dashed box).

Elsewhere on the contiguous allosteric back face of PTP1B, in the 197 site (Keedy et al., 2018), a fragment binds with a similar pose at cryo and RT (**Fig. 11**, **Fig. S12**). When this fragment binds at cryo, the protein globally remains in its default open state (**Fig. 11**). However, at RT, the allosteric L16 site closes, and the active-site WPD loop partially closes (**Fig. 11**). Notably, this fragment binds in the same position as a previously reported covalently tethered allosteric inhibitor (Keedy et al., 2018) (**Fig. S13**; see also **Fig. 9**). Thus, room temperature allows for distinct protein conformational redistributions in response to fragment binding in allosteric sites.

**Figure 11:**
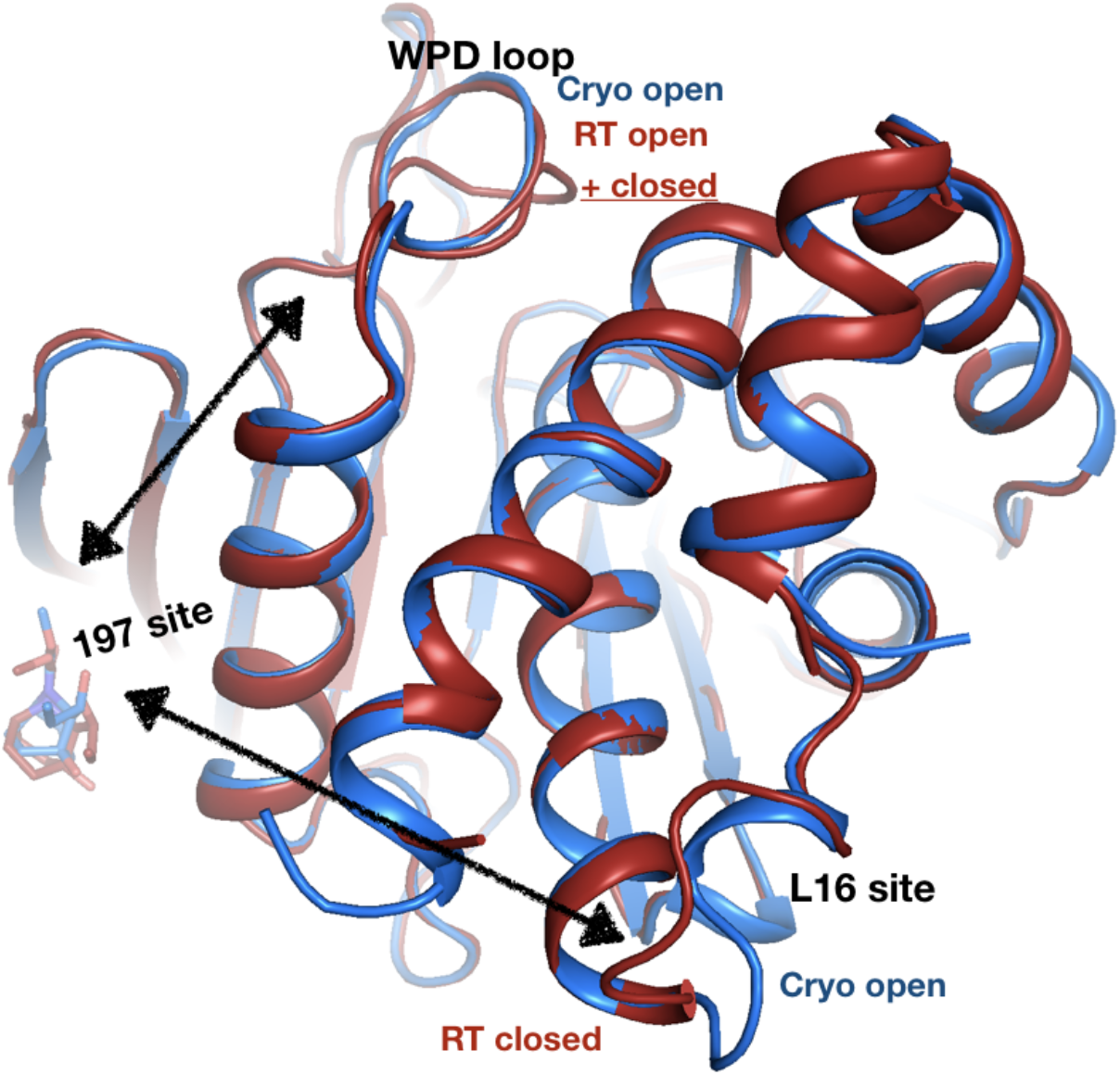
Allosteric protein responses at key sites seen only at room temperature. Although the fragment binds in a similar manner and in the same allosteric site (the 197 site) in both the RT model (z0032) (red) and the cryo model (y1763) (blue), the protein response is different between the two temperatures. At cryo, the protein retains the default open conformation, with loop 16 in the L16 site open and the WPD loop also open. Alternatively, at RT, the L16 site is fully closed, while the WPD loop exhibits alternate conformations with the loop both open and closed. The α7 helix (not shown) remains disordered in both temperatures.

## Discussion

Cryogenic X-ray crystallography is the predominant experimental method for deriving insights into protein-ligand structures, but the effects of cryo temperature on protein-ligand binding are poorly understood. To fill this critical gap, here we report a large set of room-temperature crystal structures of the dynamic enzyme PTP1B in complex with diverse small-molecule fragments, and present a detailed comparison with the corresponding cryo-temperature structures. Our data suggest that temperature can significantly affect the occupancy, pose, and even location of small-molecule binding to proteins in crystal structures. Moreover, we show that temperature can modulate protein conformational responses to ligand binding, leading to new insights into allosteric networks.

The two complementary approaches for RT crystallographic fragment screening used here led to similar conclusions, thus demonstrating the robustness of our results, but have distinct strengths. Our single-crystal approach was most directly comparable to the previous cryo experiments (Keedy et al., 2018), but is lower-throughput. By contrast, our in-situ diffraction approach was medium- to high-throughput, and can more easily minimize radiation damage, but requires merging of partial datasets that may be subtly non-isomorphous (**Fig. S2**). In this sense, the in-situ approach sits between single-crystal and serial approaches, and thus may benefit from ongoing methodological approaches in both arenas. The multi-crystal aspect of the in-situ approach poses challenges, but also opportunities: for example, future work could dissect the relative importance of clustering in reciprocal space before or after merging (Ginn, 2020) vs. averaging in real space (Pearce, Krojer, Bradley, et al., 2017) for identifying low-occupancy binding events at RT.

Although only 29-39% of the fragments that previously bound at cryo temperature (Keedy et al., 2018) also bound here at RT, several lines of evidence suggest this is predominantly due to the difference in data collection temperature, as opposed to e.g. variability in experimental steps. First, the RT hit rates for cryo-hits were similar for our two RT screens, which were performed with different techniques (single-crystal and in-situ) by largely different sets of people at different times. Second, we monitored log files from the acoustic droplet ejection instrument used for soaking (Collins et al., 2017) and excluded any crystals that may not have been soaked correctly. Third, in multiple RT datasets, a cryo-hit fragment demonstrably no longer binds at the original site but does bind at a different site (**Fig. 8**), confirming the crystals were soaked correctly. Fourth, even when cryo-hit fragments are observed in RT electron density event maps, we observe a trend of lower occupancies at RT (**Fig. 2**). We conclude that the large temperature difference between cryo and RT (>178 K) underlies the observed changes in binding. This is in accord with recent studies in which only 0 of 9 (Guven et al., 2021) and 5 of 30 (Gildea et al., 2021) cryo-hit ligands were seen to bind at RT, and in which lower occupancies were seen at RT than at cryo for <10 ligands (Bradford et al., 2021).

When fragments do bind at RT, they often do so differently than at cryo. In one example, the fragment itself has a similar pose at RT, but the surrounding solvent layer rearranges (**Fig. S8**, **Fig. 6**), akin to previous observations for proteins (Darby et al., 2019; Keedy et al., 2014; Thomaston et al., 2017); such differences in ordered solvent may be exploitable for rational inhibitor design (Balius et al., 2017; Darby et al., 2019; Ramsey et al., 2016). In a distinct type of example, the fragment binds with one pose at cryo, but a unique flipped pose at RT (**Fig. 5**). Along similar lines, another fragment binds with partial-occupancy alternate conformations (van Zundert et al., 2018) at cryo, but not at RT (**Fig. 6**). In still other examples, the fragment binds to one site at cryo, but an entirely different site at RT (**Fig. 8**, **Fig. S10**).

How does higher diffraction temperature cause such significant changes in protein-ligand binding? We speculate that after crystal soaking (which occurs at ambient temperature), all cryo-hit fragments are initially bound, but in many cases only loosely, with high B-factors that render them invisible from RT diffraction data. During cryocooling, the ligand B-factors (i.e. temperature factors) then drop rapidly on a faster timescale than overall crystal cooling (Halle, 2004), with many becoming sufficiently well-ordered to be observable in cryo density (at least with PanDDA). Relatedly, it is unclear why some fragments bind at both cryo and RT, but with a different pose or binding site: they have similar molecular weights and numbers of rotatable bonds, yet are more hydrophobic and have more interactions with the binding site (**Table S3**). To more deeply interrogate the complex relationship between temperature, cooling kinetics, and protein-ligand conformational ensembles, additional experiments are planned using mechanically controlled variable cryocooling rate (Warkentin et al., 2006) and variable crystal size. Future studies can also explore the degree to which the conclusions drawn here for small-molecule fragments can be extrapolated to larger, drug-sized ligands.

Even when ligand binding is similar at RT vs. cryo, the protein response can differ. In one case, a key secondary structural element is differentially ordered at RT vs. cryo (**Fig. 10**). In another case, the protein was inert to binding of a particular fragment at cryo, but exhibits allosteric conformational changes at multiple key sites upon binding of the same fragment in the same site at RT (**Fig. 11**). Notably, these changes involve essentially complete closing of the allosteric L16 site, but only partial closing (~50%) of the active-site WPD loop -- in contrast to the previous paradigm in which the WPD loop and allosteric sites are precisely conformationally coupled (Keedy et al., 2018). Similar (de)coupling was also seen recently with serial synchrotron crystallography of apo PTP1B (Sharma et al., 2022). Thus, RT crystallography can add important nuance to our understanding of allosteric mechanisms in PTPs (Choy et al., 2017; Cui et al., 2017; Hjortness et al., 2018) and likely other proteins.

Our results here provide several insights that can aid future development of allosteric small-molecule modulators for PTP1B, a highly validated but “undruggable” (Mullard, 2018; Zhang, 2017) therapeutic target. First, we observe new conformations for fragments on both sides of Loop 16 of the allosteric L16 site (**Fig. 5**, **Fig. 6**), offering unique footholds for structure-based inhibitor design of allosteric inhibitors. This local ligand heterogeneity, combined with the malleability of the adjacent α6 helix (**Fig. 10**, **Movie S1**) and varying levels of apparent coupling between the L16 and active sites (**Fig. 11**), argue for additional studies to decipher how different ligands in this region may selectively perturb the conformations of remote sites to allosterically control PTP1B function.

Second, one new RT fragment-binding site reported here was not previously shown to bind any fragments at cryo (Keedy et al., 2018) (although additional clustering did identify one adjacent cryo-hit (Ginn, 2020), thus offering a new ligand-binding foothold. Coincidentally, the corresponding site in the paralog SHP2 has been successfully targeted with small-molecule allosteric inhibitors (Chen et al., 2016; LaRochelle et al., 2018).

Third, we observe a fragment covalently bound to Lys197 of the allosteric 197 site, with a similar pose as our previously reported allosteric inhibitor that was covalently tethered to an engineered K197C mutant (Keedy et al., 2018) (**Fig. 9**). This unexpected result opens new doors to design a covalent allosteric inhibitor targeting wildtype PTP1B, inspired by other success stories of progressing covalent fragment hits (Miller et al., 2013; Resnick et al., 2019). The potential of targeting the allosteric 197 site of PTP1B is further reinforced by our new finding that fragment binding in this site (**Fig. S12**) can elicit allosteric conformational responses at RT that were masked at cryo (**Fig. 11**).

It is instructive to consider the results reported here in light of the growing (and exciting) trend toward leveraging artificial intelligence and machine learning to address central problems in structural biology and biophysics. Most famously, the AI/ML algorithm AlphaFold 2 (Jumper et al., 2021) (and to a lesser extent RoseTTAfold (Baek et al., 2021)) made a quantum leap in protein structure prediction accuracy. More relevant to the work reported here, AI/ML is being used to great effect for structure-based drug design and computational chemistry, including protein-ligand docking (Corso et al., 2022) and ligand design (Wallach et al., 2015). Importantly, all of these methods rely on training data in the form of experimental protein structures from the PDB, the vast majority of which are cryo-temperature crystal structures. For structure prediction, this temperature distribution undoubtedly introduces bias into the predicted models, likely favoring well-packed states that preclude functionally required conformational heterogeneity. For drug design, it may favor protein-ligand interactions that overweight enthalpic considerations and underweight entropic ones, feature inaccurate solvation environments, or suggest artificially rigid proteins. The full implications of these biases, and the prospects for bypassing or ameliorating them by other means, remain to be clarified (Bradford et al., 2021). The findings and data reported here for protein-ligand interactions at RT may provide useful tools to examine these important questions as the age of AI/ML continues to rapidly unfold.

Overall, our work highlights the value and accessibility of room-temperature crystallographic ligand screening for providing unique insights into protein-ligand interactions, particularly for allosteric sites (Krojer et al., 2020). More broadly, by using temperature as a readily accessible experimental knob, this study speaks to the potential of a multitemperature crystallography strategy, including excursions to higher temperatures in the physiological regime (Doukov et al., 2020; Ebrahim et al., 2022; Otten et al., 2020), for elucidating fundamental connections between molecular structure, heterogeneity, and function (Keedy, 2019).

## Materials and Methods

### Protein expression

All experiments used the same PTP1B construct as was used previously: residues 1-321, WT* (C32S/C92V double mutation), in the pET24b vector carrying a kanamycin resistance gene (Keedy et al., 2018). Expression and purification were also performed as previously described (Keedy et al., 2018). PTP1B was transformed into BL21 *E. coli* competent cells. The cultures were grown overnight in a 5 mL LB media containing 35 mg/L (final) kanamycin at 37°C shaking continuously at 150 rpm. Next, this overnight culture was used to inoculate 1 L LB media containing 35 mg/L (final) kanamycin. This culture was grown until the optical density at 600 nm (OD_600_) reached between 0.6-0.8. PTP1B expression was then immediately induced by adding IPTG to 100 μM (final) and incubating for about 18-20 hours at 18°C shaking continuously at 200-250 rpm. The culture was then pelleted by centrifugation, the supernatant discarded, and the cell pellets (“cellets”) harvested and stored at −80°C for subsequent purification.

### Protein purification

On the day of purification, each cellet was retrieved from −80°C and thawed on ice in 45 mL of lysis buffer (100 mM MES pH 6.5, 1 mM EDTA, freshly added 1 mM DTT) and a dissolved Pierce Protease Inhibitor Tablet. The cells were resuspended using a tabletop vortex. The homogenous cell suspension was then subjected to sonication using a Branson Digital Sonifier, with the probe submerged halfway into the suspension for about 20 minutes (10 seconds on/off) with 50% amplitude. The lysed cells were then subjected to centrifugation at 4°C, and the supernatant was filtered using 0.22 μm syringe filters and loaded onto an SP FF 16/10 cation exchange column, pre-equilibrated in lysis buffer, in an ÄKTA Pure purification system (GE Healthcare Life Sciences). The protein was eluted as 5 mL fractions using a linear gradient of lysis buffer from 0 to 1 M NaCl. PTP1B eluted at approximately 200 mM NaCl per the UV detector and SDS-PAGE analysis. The PTP1B fractions were pooled together and concentrated to 3 mL volume, then applied to a Superdex 75 (GE Healthcare Life Sciences) size exclusion column pre-equilibrated in crystallization buffer (10 mM Tris pH 7.5, 0.2 mM EDTA, 25 mM NaCl, 3 mM freshly added DTT). PTP1B eluted as a single peak, with high purity per SDS-PAGE analysis. The purified PTP1B protein was then concentrated to 40 mg/mL and used for crystallization.

### Protein crystallization

The PTP1B crystallization conditions used here were similar to those described previously (Keedy et al., 2018). 40 mg/mL protein in crystallization buffer was mixed with well solution (0.1 M HEPES pH 7.5, 0.3 M magnesium acetate, 13.5% PEG 8000, 2% ethanol, and 1 mM beta-mercaptoethanol (BME)) and seed stock in a 135:135:30 nL protein:well:seed ratio. Glycerol was not included. Seed stocks were prepared using Hampton Seed Bead tools with previously grown crystals. Drops were set using a TTP Labtech Mosquito device in 96-well sitting-drop crystallization trays. For the single-crystal screen, both MiTeGen In-Situ-1 and MRC SwissCi trays were used. For the in-situ crystallography screen, MiTeGen In-Situ-1 trays were used. Crystals appeared within about 3 days, and grew to maximum size within about 1 week. Crystals grew to dimensions of approximately 100 x 20 x 20 μm up to approximately 500 x 100 x 100 μm.

### Fragment selection

For the 1-xtal screen, we used fragments from the Maybridge 1000 fragment library (Maybridge Ro3 core set), the Edelris Keymical fragment library, and the Diamond Light Source in-house fragment library (DSPL) (Cox et al., 2016). For cryo-hits, we included 59 fragments that bound to several different sites of interest at cryo. For cryo-non-hits, we included 51 fragments that spanned the range of highly similar to dissimilar as compared to the previous cryo-hits.

For the in-situ screen, we used fragments from the DSi-Poised (DSiP) library, which is a new version of the DSPL that contains many of the same fragments. For cryo-hits, we included all cryo-hits that were available in the DSiP library, as well as 12 cryo-hits we had previously purchased, for a total of 48 molecules. For cryo-non-hits, we included the 50 fragments in the DSiP library that were most similar to any previous cryo-hit. For both screens, similarity between fragments was assessed based on Tanimoto scores calculated using RDKit (*RDKit: Open-Source Cheminformatics*, n.d.) topological fingerprints.

Some fragments that were cryo-non-hits in our original cryo screen (Keedy et al., 2018) were subsequently identified as cryo-hits using the new cluster4x method for computational clustering method (Ginn, 2020). Here, for both screens, we considered such fragments to be cryo-hits. This corresponded to 3 fragments for 1-xtal and 1 fragment for in-situ. However, no RT binding events were seen for any of these newly identified cryo-hits.

### Crystal soaking

For each screen, crystals were soaked with small-molecule fragments using an Echo acoustic droplet ejection liquid handler and a database mapping individual fragments to individual crystals, as described (Collins et al., 2017). For the in-situ screen, anywhere from 1 to 5 wells were soaked with a given fragment, depending on the number of crystals per well.

Two strategies were used to confirm that fragments were successfully soaked into the crystallization drops. First, for both screens, log files for the acoustic droplet ejection device were inspected, and any wells with suspicious entries or errors were excluded. Second, for the in-situ screen, optical images of the drops after soaking were visually inspected, and any wells that did not clearly feature a second adjacent drop corresponding to the fragment in DMSO were excluded.

### X-ray diffraction

For the 1-xtal screen, harvested crystals on size-matched nylon loops were enclosed in plastic capillaries containing ~10 μL of well solution and sealed with vacuum grease, and these samples were mounted onto the goniometer at Diamond Light Source beamline i03. Most datasets were collected with 180° of rotation over 1800 images with 0.1° oscillations with 0.05 s exposures. Some datasets near the end of the data collection shift were lowered to collect only 120° of crystal rotation, as smaller crystals sometimes did not appear to survive the full 180° dose. The X-ray beam was attenuated to 4.5% transmission for a flux of ~4.5e11 ph/s with a 50 x 20 μm or 80 x 20 μm beam profile at a wavelength of 0.97625 Å. Temperature was controlled at 278 K using an Oxford Cryostream (800 Series).

For the in-situ screen, crystallization trays were mounted onto the goniometer at Diamond Light Source beamline i24 for diffraction data collection. Partial datasets (wedges) were collected with up to 36° of rotation over 360 images with 0.1° oscillations with 0.03 s exposures. For each fragment, anywhere from 2 to 24 (average: 7) wedges were collected. In some cases, wedges for the same fragment derived from different crystals in the same well; in other cases, wedges for the same fragment derived from crystals in different wells soaked with the same fragment. The X-ray beam was attenuated to 1.5% transmission for a flux of ~4.5e10 ph/s with a 50 x 50 μm beam profile at a wavelength of 0.96874 Å. Temperature was controlled by pointing a cryostream set to 277 K at the in situ tray mounted on the goniometer. Temperature was confirmed to be ~22°C (~295 K) by a handheld thermometer held by the tray.

Translational/vector data collection was not used for either screen. Whereas cryo datasets were previously named y#### (y for “cryo”), RT datasets here were named x#### for the in-situ screen and z#### for the 1-xtal screen.

### X-ray data processing

For the 1-xtal screen, datasets were reduced using XDS (Kabsch, 2010). The frames that were used to process the datasets were manually chosen to exclude frames where the number of detected spots dipped below around 20, commonly due to the crystal rotating out of the beam, the crystal reaching the end of its lifetime due to radiation damage, or when the diffraction quality dropped as a result of the dimensions of the crystal. Multiple datasets were merged only if they derived from the same crystal. Resolution cutoffs were chosen to ensure the following statistics in the highest resolution bin: an <I/σ(I)> of 1.0 or higher, a completeness of 90% or higher, and a CC1/2 of at least 50%. The resolutions of individual datasets were not held to be identical, and the cutoff for each dataset was chosen to be the point at which the reflections from the highest resolution bin made the statistics of that bin better, or kept the same for <I/σ(I)>, CC_1/2_ and completeness. Datasets shared a common set of R_free_ flags and a common reference dataset to ensure consistent data indexing due to the space group of the crystal form, P 31 2 1. The final datasets were reasonably high-resolution (**Fig. 1A**).

For the in-situ screen, individual wedges were first reduced using Dials (Winter et al., 2018). All frames were included. Resolution cutoffs for individual wedges were chosen automatically by Dials (Winter et al., 2018). Next, multiple wedges for the same fragment, regardless of crystal or well, were merged using xia2.multiplex (Gildea et al., 2021). In some cases the unit cell (89.6, 89.6, 106.2, 90, 90, 120) and or the space group (P 31 2 1) was flagged in the xia2.multiplex input. Additionally, for some datasets the final merging step had to be done separately with dials.merge. For DMSO, anywhere from 2 to 6 wedges were merged. DMSO wedges were usually grouped by crystallization well, but in some cases were combined across wells to improve statistics. The final datasets were high-resolution (**Fig. 1B**).

To check for global radiation damage, we used RADDOSE-3D to calculate the predicted average diffraction weighted dose (ADWD) for each dataset (Bury et al., 2018). For the in-situ screen, predicted ADWD was ~0.03-0.04 MGy, depending on estimated crystal size, using the up-to-36° wedges. For the 1-xtal screen, predicted ADWD was ~3.2-7.4 MGy, depending on estimated crystal size and beam size, using the full 180° datasets. Thus the in-situ data are well below the estimated RT limit of ~0.4 MGy (Fischer, 2021). The 1-xtal data are above the quoted RT limit (yet below the cryo limit of ~20-30 MGy (Owen et al., 2006)); this was ameliorated for individual datasets by cutting later frames with reduced average intensities, as noted above. Additionally, we inspected 2Fo-Fc electron density maps for the individual 1-xtal RT hits featured here and observed no signatures of local radiation damage such as decarboxylation of Asp/Glu sidechains, whether near the cryo and/or RT fragment binding site(s) or elsewhere in the protein.

For an alternative data processing pipeline for the in-situ data, the cluster4x algorithm (Ginn, 2020) was used to pre-cluster in-situ wedges before merging with xia2.multiplex. First, the P 31 2 1 indexing hand for approximately half the wedges was changed using the Pointless (Evans, 2006) utility from CCP4 (Winn et al., 2011) to achieve consistency. Then, the wedges were clustered in real space. The resulting three clusters were partially overlapping in this space, and datasets were visually/manually assigned to these clusters. For each cluster, xia2.multiplex was used as described above, and separate PanDDa runs were performed as described below.

For each dataset, we used the Dimple utility from CCP4 (Winn et al., 2011) for phasing and initial refinement. Dimple was run with molecular replacement (flag: -M0) for the first dataset only, and only with downstream refinement steps (flag: -M1) for all other datasets. Additional flags were included to obtain a consistent set of R_free_ reflections (--free-r-flags, --freecolumn R-free-flags). For both screens, the same structural model was used for Dimple, based on a high-resolution DMSO-soaked in-situ merged dataset. This model reflects the predominant global open state of PTP1B, with the α7 helix unmodeled and the C-terminus of the α6 helix modeled with partial occupancy (Keedy et al., 2018).

### PanDDA modeling and refinement

For both the 1-xtal and in-situ screens, PanDDA (Pearce, Krojer, Bradley, et al., 2017) version 0.2.14 was used. The pandda.analyse command was used with the minimum build datasets set to 20.

In addition to the automatic PanDDA analysis, for each dataset for which PanDDA did not show an event, we did a manual BDC scan from 1-BDC values of 0 to 0.9 as well as generating custom maps at the 1-BDC value that corresponded to the cryo 1-BDC. We saw 5 events with this manual inspection that PanDDA missed at the corresponding cryo 1-BDC. We used the automatically generated event maps throughout the manuscript, unless otherwise noted that a manually calculated event map is used.

Fragments and associated protein changes were modeled using pandda.inspect in Coot. Waters were kept the same between the unbound and bound models, except where the PanDDA event map indicated a shift, deletion, or an addition of a new water. Ligand restraints files were calculated with eLBOW (Moriarty et al., 2009). We aimed to keep the RT models similar to the cryo models except when the RT map argued otherwise, so that modeled differences were due to temperature.

For the in-situ datasets in this manuscript, we report all hits derived from the all-wedges datasets, plus a small number of distinct hits from the pre-clustered datasets as noted where appropriate.

Because ligands are not fully occupied, to prepare for refinement we must use an ensemble of bound state plus unbound i.e. ground state for refinement (Pearce, Krojer, & von Delft, 2017). We generated such an ensemble model with pandda.export. We then added hydrogens with Phenix ReadySet! Restraints, both between multi state occupancy groups and between local alternate locations, were generated using giant.make_restraint scripts from PanDDA 1.0.0. The argument ‘MAKE HOUT Yes’ was added to the Refmac restraint file to ensure the Hydrogens were preserved.

For refinement of fragment-bound ensemble models, the published protocol for post-PanDDA refinement for deposition (Pearce, Krojer, & von Delft, 2017) was used, including the giant.quick_refine scripts from PanDDA 1.0.0 and the program Refmac (Murshudov et al., 2011). For a few examples, the script was rerun if the ligand was refined to a total occupancy greater than 1. Additionally, some hydrogens refined to 0 occupancy so they were manually edited to match the remainder of its residue. Refined bound-state models were then re-extracted using giant.split_conformations.

In addition to fragment-bound models, a ground-state model was refined for each screen, using the model used for MR previously and the highest-resolution DMSO dataset per screen.

## Data availability

Bound state-models, structure factors, PanDDA event maps, and traditional maps (2Fo-Fc and Fo-Fc) for all fragment-bound structures are available in the Protein Data Bank under the following PDB ID accession codes: 7FQM, 7FQN, 7FQO, 7FQP, 7FQQ, 7FQR, 7FQS, 7FQT, 7FQU, 7FQV, 7FQW, 7FQX, 7FQY, 7FQZ, 7FRF, 7FRG, 7FRH, 7FRI, 7FRJ, 7FRK, 7FRL, 7FRM, 7FRN, 7FRO, 7FRP, 7FRQ, 7FRR. For each screen, a ground-state (unbound) model is also available, along with structure factors for all datasets involved in the respective screen, under the following PDB ID accession codes: 7FRE (1-xtal), 7FRS (in-situ), 7FRT (in-situ, cluster 1), 7FRU (in-situ, cluster 2).

In addition, we provide a Zenodo directory containing our full PanDDA run directories, bound-state models, event maps, identifying information for all fragments, and related details at the following DOI: 10.5281/zenodo.7255364.

## Funding & Acknowledgements

TS was the recipient of a fellowship award from the U.S. Department of Education Graduate Assistance in Areas of National Need (GAANN) Program in Molecular Biophysics and Biomaterials at The City College of New York (PA200A150068), and is supported by CUNY Graduate Center Dissertation Fellowship.

JTB was supported by an NSF GRFP award.

DAK is supported by NIH R35 GM133769.

We thank James Fraser for guidance and discussions; the beamline staff for help operating the XChem fragment-screening pipeline at Diamond Light Source; George Meigs, James Holton, James Sandy, and Juan Sanchez-Weatherby for help with initial in-situ trial experiments; Helen Ginn for helpful discussions about diffraction dataset clustering; Virgil Woods and Nathanael Singh for help with a PTP1B activity assay; and Zachary Hill, Neel Shah, and Jack Taunton for helpful discussions about covalent fragment binding.

## References

Baek, M., DiMaio, F., Anishchenko, I., Dauparas, J., Ovchinnikov, S., Lee, G. R., Wang, J., Cong, Q., Kinch, L. N., Schaeffer, R. D., Millán, C., Park, H., Adams, C., Glassman, C. R., DeGiovanni, A., Pereira, J. H., Rodrigues, A. V., van Dijk, A. A., Ebrecht, A. C.,… Baker, D. (2021). Accurate prediction of protein structures and interactions using a three-track neural network. Science, 373(6557), 871–876.

Balius, T. E., Fischer, M., Stein, R. M., Adler, T. B., Nguyen, C. N., Cruz, A., Gilson, M. K., Kurtzman, T., & Shoichet, B. K. (2017). Testing inhomogeneous solvation theory in structure-based ligand discovery. Proceedings of the National Academy of Sciences of the United States of America, 114(33), E6839–E6846.

Berman, H. M., Westbrook, J., Feng, Z., Gilliland, G., Bhat, T. N., Weissig, H., Shindyalov, I. N., & Bourne, P. E. (2000). The Protein Data Bank. Nucleic Acids Research, 28(1), 235–242.

Bradford, S. Y. C., El Khoury, L., Ge, Y., Osato, M., Mobley, D. L., & Fischer, M. (2021). Temperature artifacts in protein structures bias ligand-binding predictions. Chemical Science. https://doi.org/10.1039/D1SC02751D

Bury, C. S., Brooks-Bartlett, J. C., Walsh, S. P., & Garman, E. F. (2018). Estimate your dose:RADDOSE-3D. Protein Science: A Publication of the Protein Society, 27(1), 217–228.

Chen, Y.-N. P., LaMarche, M. J., Chan, H. M., Fekkes, P., Garcia-Fortanet, J., Acker, M. G., Antonakos, B., Chen, C. H.-T., Chen, Z., Cooke, V. G., Dobson, J. R., Deng, Z., Fei, F., Firestone, B., Fodor, M., Fridrich, C., Gao, H., Grunenfelder, D., Hao, H.-X.,… Fortin, P. D. (2016). Allosteric inhibition of SHP2 phosphatase inhibits cancers driven by receptor tyrosine kinases. Nature, 535(7610), 148–152.

Choy, M. S., Li, Y., Machado, L. E. S. F., Kunze, M. B. A., Connors, C. R., Wei, X., Lindorff-Larsen, K., Page, R., & Peti, W. (2017). Conformational Rigidity and Protein Dynamics at Distinct Timescales Regulate PTP1B Activity and Allostery. Molecular Cell, 65(4), 644–658.e5.

Collins, P. M., Ng, J. T., Talon, R., Nekrosiute, K., Krojer, T., Douangamath, A., Brandao-Neto, J., Wright, N., Pearce, N. M., & von Delft, F. (2017). Gentle, fast and effective crystal soaking by acoustic dispensing. Acta Crystallographica. Section D, Structural Biology, 73(Pt 3), 246–255.

Corso, G., Stärk, H., Jing, B., Barzilay, R., & Jaakkola, T. (2022). DiffDock: Diffusion Steps, Twists,and Turns for Molecular Docking. In arXiv [q-bio.BM]. arXiv. http://arxiv.org/abs/2210.01776

Cox, O. B., Krojer, T., Collins, P., Monteiro, O., Talon, R., Bradley, A., Fedorov, O., Amin, J., Marsden, B. D., Spencer, J., & Others. (2016). A poised fragment library enables rapid synthetic expansion yielding the first reported inhibitors of PHIP (2), an atypical bromodomain. Chemical Science, 7(3), 2322–2330.

Cui, D. S., Beaumont, V., Ginther, P. S., Lipchock, J. M., & Loria, J. P. (2017). Leveraging Reciprocity to Identify and Characterize Unknown Allosteric Sites in Protein Tyrosine Phosphatases. Journal of Molecular Biology, 429(15), 2360–2372.

Daina, A., Michielin, O., & Zoete, V. (2017). SwissADME: a free web tool to evaluate pharmacokinetics, drug-likeness and medicinal chemistry friendliness of small molecules. Scientific Reports, 7, 42717.

Darby, J. F., Hopkins, A. P., Shimizu, S., Roberts, S. M., Brannigan, J. A., Turkenburg, J. P., Thomas, G. H., Hubbard, R. E., & Fischer, M. (2019). Water Networks Can Determine the Affinity of Ligand Binding to Proteins. Journal of the American Chemical Society, 141(40), 15818–15826.

Douangamath, A., Fearon, D., Gehrtz, P., Krojer, T., Lukacik, P., Owen, C. D., Resnick, E., Strain-Damerell, C., Aimon, A., Ábrányi-Balogh, P., Brandão-Neto, J., Carbery, A., Davison, G., Dias, A., Downes, T. D., Dunnett, L., Fairhead, M., Firth, J. D., Jones, S. P.,… Walsh, M. A. (2020). Crystallographic and electrophilic fragment screening of the SARS-CoV-2 main protease. Nature Communications, 11(1), 5047.

Doukov, T., Herschlag, D., & Yabukarski, F. (2020). Instrumentation and experimental procedures for robust collection of X-ray diffraction data from protein crystals across physiological temperatures. Journal of Applied Crystallography, 53(Pt 6), 1493–1501.

Ebrahim, A., Riley, B. T., Kumaran, D., Andi, B., Fuchs, M. R., McSweeney, S., & Keedy, D. A. (2022). The temperature-dependent conformational ensemble of SARS-CoV-2 main protease (Mpro). IUCrJ, 9(5). https://doi.org/10.1107/S2052252522007497

Elchebly, M., Payette, P., Michaliszyn, E., Cromlish, W., Collins, S., Loy, A. L., Normandin, D., Cheng, A., Himms-Hagen, J., Chan, C. C., Ramachandran, C., Gresser, M. J., Tremblay, M. L., & Kennedy, B. P. (1999). Increased insulin sensitivity and obesity resistance in mice lacking the protein tyrosine phosphatase-1B gene [Review of *Increased insulin sensitivity and obesity resistance in mice lacking the protein tyrosine phosphatase-1B gene*]. Science, 283(5407), 1544–1548.

Evans, P. (2006). Scaling and assessment of data quality. Acta Crystallographica. Section D,Biological Crystallography, 62(Pt 1), 72–82.

Fenwick, R. B., van den Bedem, H., Fraser, J. S., & Wright, P. E. (2014). Integrated description of protein dynamics from room-temperature X-ray crystallography and NMR. Proceedings of the National Academy of Sciences of the United States of America, 111(4), E445–E454.

Fischer, M. (2021). Macromolecular room temperature crystallography. Quarterly Reviews of Biophysics, 54, e1.

Fischer, M., Shoichet, B. K., & Fraser, J. S. (2015). One Crystal, Two Temperatures: Cryocooling Penalties Alter Ligand Binding to Transient Protein Sites. Chembiochem: A European Journal of Chemical Biology, 16(11), 1560–1564.

Fraser, J. S., Clarkson, M. W., Degnan, S. C., Erion, R., Kern, D., & Alber, T. (2009). Hidden alternative structures of proline isomerase essential for catalysis. Nature, 462(7273), 669–673.

Fraser, J. S., van den Bedem, H., Samelson, A. J., Lang, P. T., Holton, J. M., Echols, N., & Alber, T. (2011). Accessing protein conformational ensembles using room-temperature X-ray crystallography. Proceedings of the National Academy of Sciences of the United States of America, 108(39), 16247–16252.

Gelin, M., Delfosse, V., Allemand, F., Hoh, F., Sallaz-Damaz, Y., Pirocchi, M., Bourguet, W., Ferrer, J.-L., Labesse, G., & Guichou, J.-F. (2015). Combining ‘dry’ co-crystallization and in situ diffraction to facilitate ligand screening by X-ray crystallography. Acta Crystallographica. Section D, Biological Crystallography, 71(8), 1777–1787.

Gildea, R. J., Beilsten-Edmands, J., Axford, D., Horrell, S., Owen, R. L., Winter, G., & IUCr. (2021). xia2.multiplex: a multi-crystal data analysis pipeline. Acta Crystallographica Section A: Foundations and Advances, 77, C149–C149.

Ginn, H. M. (2020). Pre-clustering data sets using cluster4x improves the signal-to-noise ratio of high-throughput crystallography drug-screening analysis. Acta Crystallographica. Section D, Structural Biology, 76(Pt 11), 1134–1144.

Günther, S., Reinke, P. Y. A., Fernández-García, Y., Lieske, J., Lane, T. J., Ginn, H. M., Koua, F. H. M., Ehrt, C., Ewert, W., Oberthuer, D., Yefanov, O., Meier, S., Lorenzen, K., Krichel, B., Kopicki, J.-D., Gelisio, L., Brehm, W., Dunkel, I., Seychell, B.,… Meents, A. (2021). X-ray screening identifies active site and allosteric inhibitors of SARS-CoV-2 main protease. Science, 372(6542), 642–646.

Guven, O., Gul, M., Ayan, E., Johnson, J. A., Cakilkaya, B., Usta, G., Ertem, F. B., Tokay, N., Yuksel, B., Gocenler, O., Buyukdag, C., Botha, S., Ketawala, G., Su, Z., Hayes, B., Poitevin, F., Batyuk, A., Yoon, C. H., Kupitz, C.,… DeMirci, H. (2021). Case Study of High-Throughput Drug Screening and Remote Data Collection for SARS-CoV-2 Main Protease by Using Serial Femtosecond X-ray Crystallography. Crystals, 11(12), 1579.

Halle, B. (2004). Biomolecular cryocrystallography: structural changes during flash-cooling. Proceedings of the National Academy of Sciences of the United States of America, 101(14), 4793–4798.

Hjortness, M. K., Riccardi, L., Hongdusit, A., Zwart, P. H., Sankaran, B., De Vivo, M., & Fox, J. M. (2018). Evolutionarily Conserved Allosteric Communication in Protein Tyrosine Phosphatases. Biochemistry, 57(45), 6443–6451.

Hongdusit, A., Zwart, P. H., Sankaran, B., & Fox, J. M. (2020). Minimally disruptive optical control of protein tyrosine phosphatase 1B. Nature Communications, 11(1), 788.

Jumper, J., Evans, R., Pritzel, A., Green, T., Figurnov, M., Ronneberger, O., Tunyasuvunakool, K., Bates, R., Žídek, A., Potapenko, A., Bridgland, A., Meyer, C., Kohl, S. A. A., Ballard, A. J., Cowie, A., Romera-Paredes, B., Nikolov, S., Jain, R., Adler, J.,… Hassabis, D. (2021). Highly accurate protein structure prediction with AlphaFold. Nature, 596(7873), 583–589.

Kabsch, W. (2010). XDS. Acta Crystallographica. Section D, Biological Crystallography, 66(Pt 2), 125–132.

Keedy, D. A. (2019). Journey to the center of the protein: allostery from multitemperature multiconformer X-ray crystallography. Acta Crystallographica. Section D, Structural Biology, 75(Pt 2), 123–137.

Keedy, D. A., Hill, Z. B., Biel, J. T., Kang, E., & Rettenmaier, T. J. (2018). An expanded allosteric network in PTP1B by multitemperature crystallography, fragment screening, and covalent tethering. eLife. https://elifesciences.org/articles/36307

Keedy, D. A., Kenner, L. R., Warkentin, M., Woldeyes, R. A., Hopkins, J. B., Thompson, M. C., Brewster, A. S., Van Benschoten, A. H., Baxter, E. L., Uervirojnangkoorn, M., McPhillips, S. E., Song, J., Alonso-Mori, R., Holton, J. M., Weis, W. I., Brunger, A. T., Soltis, S. M., Lemke, H., Gonzalez, A.,… Fraser, J. S. (2015). Mapping the conformational landscape of a dynamic enzyme by multitemperature and XFEL crystallography. eLife, 4. https://doi.org/10.7554/eLife.07574

Keedy, D. A., Van Den Bedem, H., Sivak, D. A., & Petsko, G. A. (2014). Crystal cryocooling distorts conformational heterogeneity in a model Michaelis complex of DHFR. Structure. https://www.sciencedirect.com/science/article/pii/S0969212614001403

Krishnan, N., Koveal, D., Miller, D. H., Xue, B., Akshinthala, S. D., Kragelj, J., Jensen, M. R., Gauss, C.-M., Page, R., Blackledge, M., Muthuswamy, S. K., Peti, W., & Tonks, N. K. (2014). Targeting the disordered C terminus of PTP1B with an allosteric inhibitor. Nature Chemical Biology, 10(7), 558–566.

Krishnan, N., Krishnan, K., Connors, C. R., Choy, M. S., Page, R., Peti, W., Van Aelst, L., Shea, S. D., & Tonks, N. K. (2015). PTP1B inhibition suggests a therapeutic strategy for Rett syndrome. The Journal of Clinical Investigation, 125(8), 3163–3177.

Krojer, T., Fraser, J. S., & von Delft, F. (2020). Discovery of allosteric binding sites by crystallographic fragment screening. Current Opinion in Structural Biology, 65, 209–216.

LaRochelle, J. R., Fodor, M., Vemulapalli, V., Mohseni, M., Wang, P., Stams, T., LaMarche, M. J., Chopra, R., Acker, M. G., & Blacklow, S. C. (2018). Structural reorganization of SHP2 by oncogenic mutations and implications for oncoprotein resistance to allosteric inhibition. Nature Communications, 9(1), 4508.

Lieske, J., Cerv, M., Kreida, S., Komadina, D., Fischer, J., Barthelmess, M., Fischer, P., Pakendorf, T., Yefanov, O., Mariani, V., Seine, T., Ross, B. H., Crosas, E., Lorbeer, O., Burkhardt, A., Lane, T. J.,Guenther, S., Bergtholdt, J., Schoen, S.,… Meents, A. (2019). On-chip crystallization for serial crystallography experiments and on-chip ligand-binding studies. IUCrJ, 6(Pt 4), 714–728.

Maeki, M., Ito, S., Takeda, R., Ueno, G., Ishida, A., Tani, H., Yamamoto, M., & Tokeshi, M. (2020). Room-temperature crystallography using a microfluidic protein crystal array device and its application to protein-ligand complex structure analysis. Chemical Science, 11(34), 9072–9087.

Milano, S. K., Huang, Q., Nguyen, T.-T. T., Ramachandran, S., Finke, A., Kriksunov, I., Schuller, D. J., Szebenyi, D. M., Arenholz, E., McDermott, L. A., Sukumar, N., Cerione, R. A., & Katt, W. P. (2022). New insights into the molecular mechanisms of glutaminase C inhibitors in cancer cells using serial room temperature crystallography. The Journal of Biological Chemistry, 298(2), 101535.

Miller, R. M., Paavilainen, V. O., Krishnan, S., Serafimova, I. M., & Taunton, J. (2013). Electrophilic fragment-based design of reversible covalent kinase inhibitors. Journal of the American Chemical Society, 135(14), 5298–5301.

Moreno-Chicano, T., Ebrahim, A., Axford, D., Appleby, M. V., Beale, J. H., Chaplin, A. K., Duyvesteyn, H. M. E., Ghiladi, R. A., Owada, S., Sherrell, D. A., Strange, R. W., Sugimoto, H., Tono, K., Worrall, J. A. R., Owen, R. L., & Hough, M. A. (2019). High-throughput structures of protein–ligand complexes at room temperature using serial femtosecond crystallography. IUCrJ, 6(6), 1074–1085.

Moriarty, N. W., Grosse-Kunstleve, R. W., & Adams, P. D. (2009). electronic Ligand Builder and Optimization Workbench (eLBOW): a tool for ligand coordinate and restraint generation. Acta Crystallographica. Section D, Biological Crystallography, 65(Pt 10), 1074–1080.

Mullard, A. (2018). Phosphatases start shedding their stigma of undruggability. Nature Reviews. Drug Discovery, 17(12), 847–849.

Murshudov, G. N., Skubák, P., Lebedev, A. A., Pannu, N. S., Steiner, R. A., Nicholls, R. A., Winn, M. D., Long, F., & Vagin, A. A. (2011). REFMAC5 for the refinement of macromolecular crystal structures. Acta Crystallographica. Section D, Biological Crystallography, 67(Pt 4), 355–367.

Otten, R., Pádua, R. A. P., Bunzel, H. A., Nguyen, V., Pitsawong, W., Patterson, M., Sui, S., Perry, S.L., Cohen, A. E., Hilvert, D., & Kern, D. (2020). How directed evolution reshapes the energy landscape in an enzyme to boost catalysis. Science, 370(6523), 1442–1446.

Owen, R. L., Rudiño-Piñera, E., & Garman, E. F. (2006). Experimental determination of the radiation dose limit for cryocooled protein crystals. Proceedings of the National Academy of Sciences of the United States of America, 103(13), 4912–4917.

Pearce, N. M., Krojer, T., Bradley, A. R., Collins, P., Nowak, R. P., Talon, R., Marsden, B. D., Kelm, S., Shi, J., Deane, C. M., & von Delft, F. (2017). A multi-crystal method for extracting obscured crystallographic states from conventionally uninterpretable electron density. Nature Communications, 8, 15123.

Pearce, N. M., Krojer, T., & von Delft, F. (2017). Proper modelling of ligand binding requires an ensemble of bound and unbound states. Acta Crystallographica. Section D, Structural Biology, 73(Pt 3), 256–266.

Ramsey, S., Nguyen, C., Salomon-Ferrer, R., Walker, R. C., Gilson, M. K., & Kurtzman, T. (2016). Solvation thermodynamic mapping of molecular surfaces in AmberTools: GIST. Journal of Computational Chemistry, 37(21), 2029–2037.

DKit: Open-source cheminformatics. Retrieved August 18, 2022, from http://www.rdkit.org

Resnick, E., Bradley, A., Gan, J., Douangamath, A., Krojer, T., Sethi, R., Geurink, P. P., Aimon, A., Amitai, G., Bellini, D., Bennett, J., Fairhead, M., Fedorov, O., Gabizon, R., Gan, J., Guo, J., Plotnikov, A., Reznik, N., Ruda, G. F.,… London, N. (2019). Rapid Covalent-Probe Discovery by Electrophile-Fragment Screening. Journal of the American Chemical Society, 141(22), 8951–8968.

Sanchez-Weatherby, J., Sandy, J., Mikolajek, H., Lobley, C. M. C., Mazzorana, M., Kelly, J., Preece, G., Littlewood, R., & Sørensen, T. L. M. (2019). VMXi: a fully automated, fully remote, high-flux in situ macromolecular crystallography beamline. Journal of Synchrotron Radiation, 26(Pt 1), 291–301.

Sharma, S., Ebrahim, A., & Keedy, D. A. (2022). Room-temperature serial synchrotron crystallography of apo PTP1B. In bioRxiv (p. 2022.07.28.501725). https://doi.org/10.1101/2022.07.28.501725

Sui, S., Mulichak, A., Kulathila, R., McGee, J., Filiatreault, D., Saha, S., Cohen, A., Song, J., Hung, H., Selway, J., Kirby, C., Shrestha, O. K., Weihofen, W., Fodor, M., Xu, M., Chopra, R., & Perry, S. L. (2021). A capillary-based microfluidic device enables primary high-throughput room-temperature crystallographic screening. Journal of Applied Crystallography, 54(Pt 4), 1034–1046.

Teplitsky, E., Joshi, K., Ericson, D. L., Scalia, A., Mullen, J. D., Sweet, R. M., & Soares, A. S. (2015). High throughput screening using acoustic droplet ejection to combine protein crystals and chemical libraries on crystallization plates at high density. Journal of Structural Biology, 191(1), 49–58.

The COVID Moonshot Consortium, Achdout, H., Aimon, A., Bar-David, E., Barr, H., Ben-Shmuel, A., Bennett, J., Bilenko, V. A., Boby, M. L., Borden, B., Bowman, G. R., Brun, J., Bvnbs, S., Calmiano, M., Carbery, A., Carney, D., Cattermole, E., Chang, E., Chernyshenko, E.,…Zitzmann, N. (2022). Open Science Discovery of Oral Non-Covalent SARS-CoV-2 Main Protease Inhibitor Therapeutics. In bioRxiv (p. 2020.10.29.339317).https://doi.org/10.1101/2020.10.29.339317

Thomaston, J. L., Woldeyes, R. A., Nakane, T., Yamashita, A., Tanaka, T., Koiwai, K., Brewster, A. S., Barad, B. A., Chen, Y., Lemmin, T., Uervirojnangkoorn, M., Arima, T., Kobayashi, J., Masuda, T., Suzuki, M., Sugahara, M., Sauter, N. K., Tanaka, R., Nureki, O.,… DeGrado, W. F. (2017). XFEL structures of the influenza M2 proton channel: Room temperature water networks and insights into proton conduction. Proceedings of the National Academy of Sciences of the United States of America, 114(51), 13357–13362.

Torgeson, K. R., Clarkson, M. W., Granata, D., Lindorff-Larsen, K., Page, R., & Peti, W. (2022). Conserved conformational dynamics determine enzyme activity. Science Advances, 8(31), eabo5546.

van Montfort, R. L. M., Congreve, M., Tisi, D., Carr, R., & Jhoti, H. (2003). Oxidation state of the active-site cysteine in protein tyrosine phosphatase 1B. Nature, 423(6941), 773–777.

van Zundert, G. C. P., Hudson, B. M., de Oliveira, S. H. P., Keedy, D. A., Fonseca, R., Heliou, A., Suresh, P., Borrelli, K., Day, T., Fraser, J. S., & van den Bedem, H. (2018). qFit-ligand Reveals Widespread Conformational Heterogeneity of Drug-Like Molecules in X-Ray Electron Density Maps. Journal of Medicinal Chemistry, 61(24), 11183–11198.

Wallach, I., Dzamba, M., & Heifets, A. (2015). AtomNet: A Deep Convolutional Neural Network for Bioactivity Prediction in Structure-based Drug Discovery. In arXiv [cs.LG]. arXiv. http://arxiv.org/abs/1510.02855

Warkentin, M., Berejnov, V., Husseini, N. S., & Thorne, R. E. (2006). Hyperquenching for protein cryocrystallography. Journal of Applied Crystallography, 39(6), 805–811.

Weinert, T., Olieric, N., Cheng, R., Brünle, S., James, D., Ozerov, D., Gashi, D., Vera, L., Marsh, M., Jaeger, K., Dworkowski, F., Panepucci, E., Basu, S., Skopintsev, P., Doré, A. S., Geng, T., Cooke, R. M., Liang, M., Prota, A. E.,… Standfuss, J. (2017). Serial millisecond crystallography for routine room-temperature structure determination at synchrotrons. Nature Communications, 8(1), 542.

Wiesmann, C., Barr, K. J., Kung, J., Zhu, J., Erlanson, D. A., Shen, W., Fahr, B. J., Zhong, M., Taylor, L., Randal, M., McDowell, R. S., & Hansen, S. K. (2004). Allosteric inhibition of protein tyrosine phosphatase 1B. Nature Structural & Molecular Biology, 11(8), 730–737.

Winn, M. D., Ballard, C. C., Cowtan, K. D., Dodson, E. J., Emsley, P., Evans, P. R., Keegan, R. M., Krissinel, E. B., Leslie, A. G. W., McCoy, A., McNicholas, S. J., Murshudov, G. N., Pannu, N. S., Potterton, E. A., Powell, H. R., Read, R. J., Vagin, A., & Wilson, K. S. (2011). Overview of the CCP4 suite and current developments. Acta Crystallographica. Section D, Biological Crystallography, 67(Pt 4), 235–242.

Winter, G., Waterman, D. G., Parkhurst, J. M., Brewster, A. S., Gildea, R. J., Gerstel, M., Fuentes-Montero, L., Vollmar, M., Michels-Clark, T., Young, I. D., Sauter, N. K., & Evans, G. (2018). DIALS: implementation and evaluation of a new integration package. Acta Crystallographica. Section D, Structural Biology, 74(Pt 2), 85–97.

Word, J. M., Lovell, S. C., LaBean, T. H., Taylor, H. C., Zalis, M. E., Presley, B. K., Richardson, J. S., & Richardson, D. C. (1999). Visualizing and quantifying molecular goodness-of-fit: small-probe contact dots with explicit hydrogen atoms. Journal of Molecular Biology, 285(4), 1711–1733.

Zhang, Z.-Y. (2017). Drugging the Undruggable: Therapeutic Potential of Targeting Protein Tyrosine Phosphatases. Accounts of Chemical Research, 50(1), 122–129.

