## Supplemental Figures and Tables for "Room-temperature crystallography reveals altered binding of small-molecule fragments to PTP1B"

### Supplementary Information

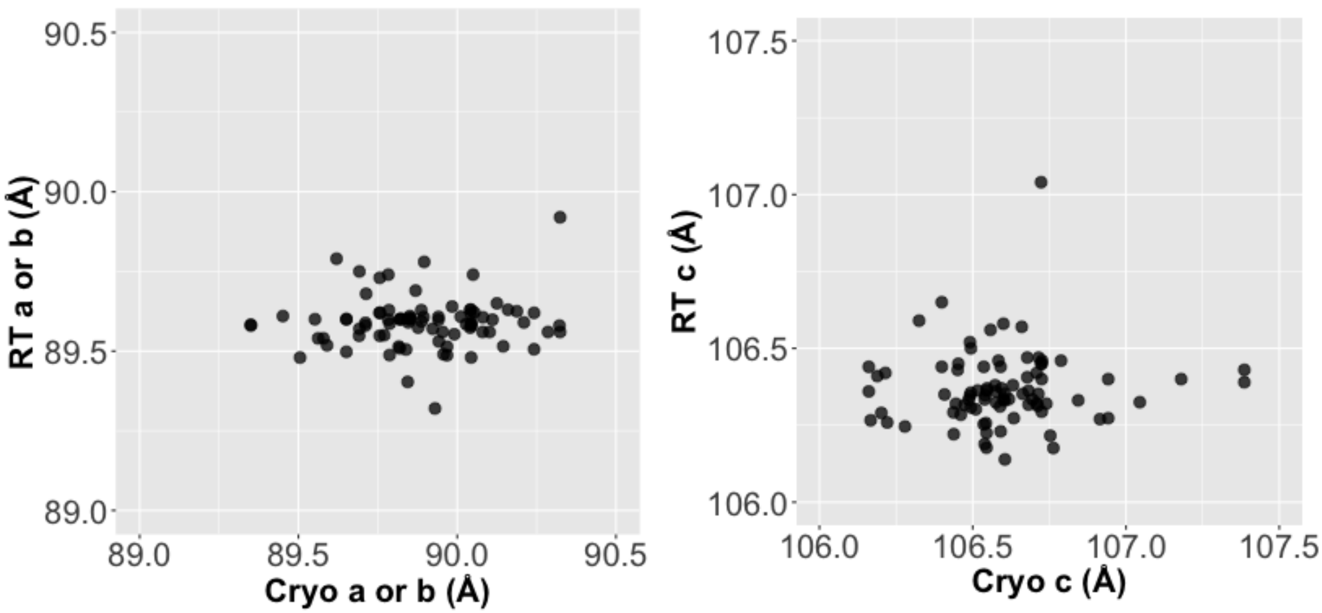


**Figure S1: Unit cell is more variable at cryo than at RT.** A comparison of RT vs cryo unit cells (*left:* cryo a or b length vs RT a or b length, *right:* cryo c length vs RT c length) reveals the cryo datasets' unit cell size has more variability than those at RT.

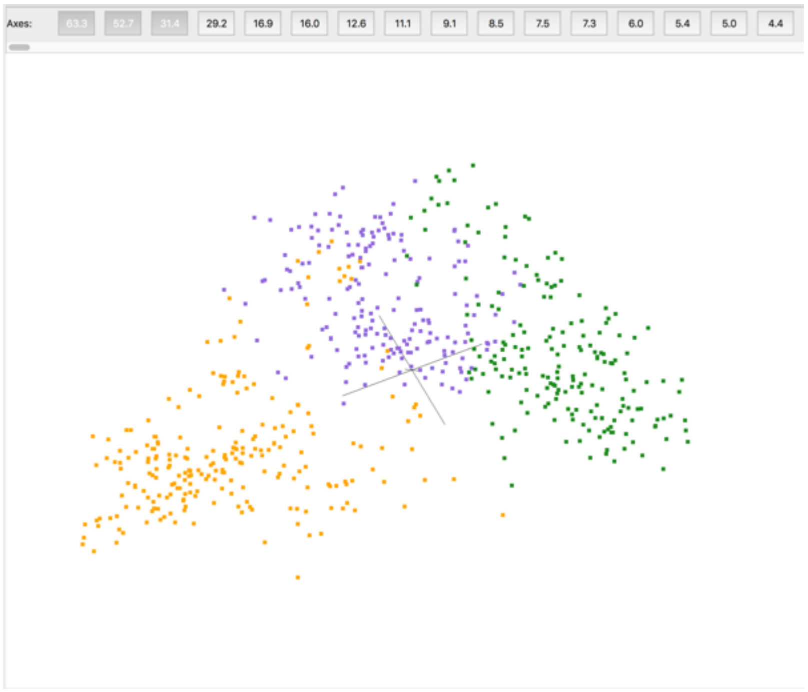


**Figure S2: Pre-clustering partial datasets from in-situ crystallography.** Partial datasets or "wedges" are represented in a reduced dimensionality space based upon structure factor amplitude differences. These wedges can be approximately divided into three clusters (orange, purple, green). Some wedges at the cluster interfaces were used in multiple clusters. Wedges within a cluster were subsequently merged to generate complete datasets for input to PanDDA. Image from cluster4x (Ginn, 2020).

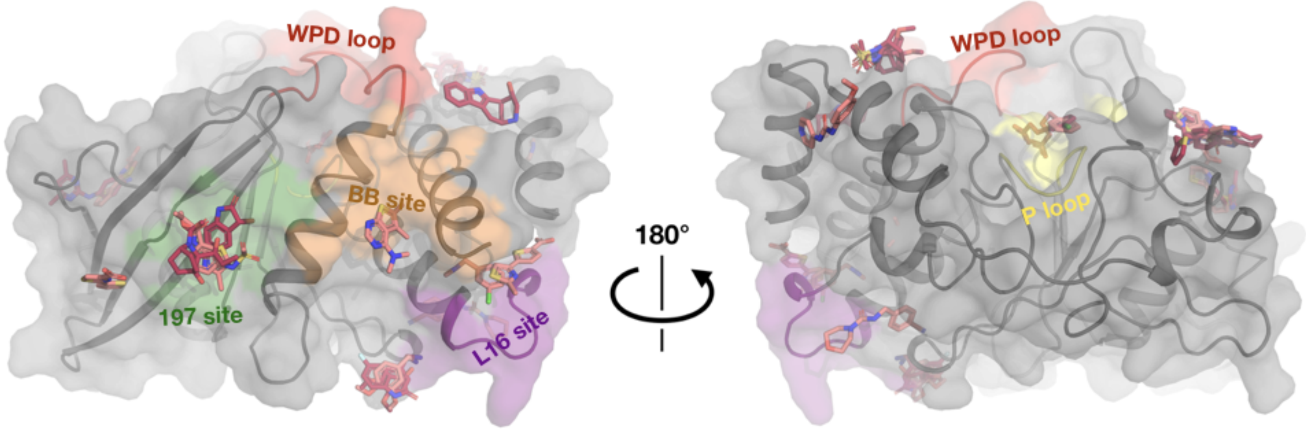


**Figure S3: Fragments bound at room temperature colored by RT screen.** Same views and fragments as **Fig. 3**, but with fragments colored based on the RT screen: 1-xtal (dark pink) or in-situ (light pink).

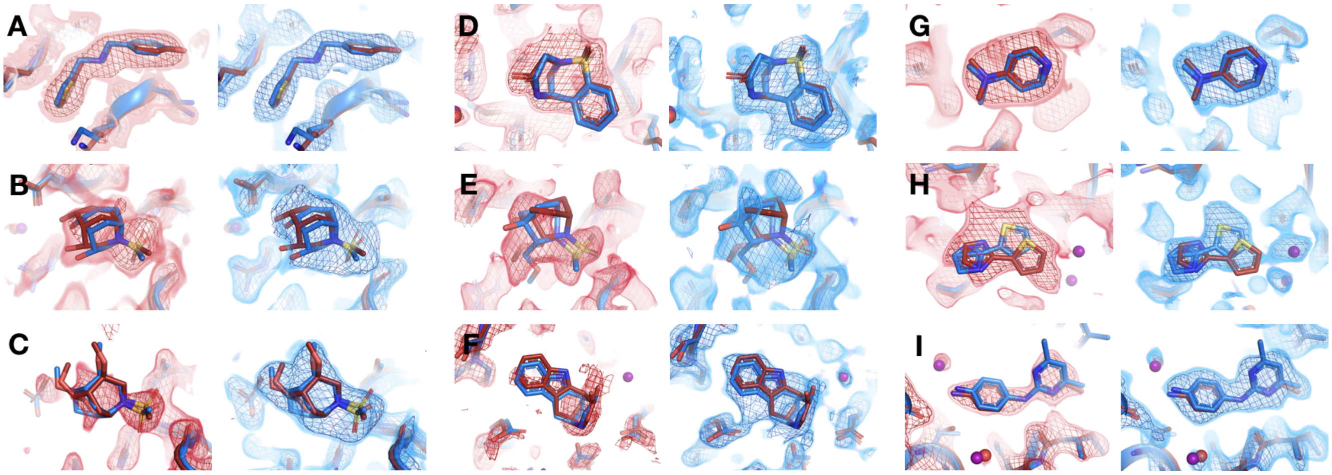


**Figure S4: All fragments that bind similarly at room vs. cryo temperatures from 1-xtal screen.** For each panel: RT event map density on left, cryo event map density on right (blue). Same coloring as main **Fig. 4.**

**(A)** RT: z0007 (2 σ), cryo: y1710 (1.2 σ).
**(B)** RT: z0015 (1.8 σ), cryo: y1554 (1.5 σ)
**(C)** RT: z0021 (1.1 σ), cryo: y1312 (1.5 σ)
**(D)** RT: z0025 (1.5 σ), cryo: y1294 (1.5 σ)
**(E)** RT: z0028 (1.2 σ), cryo: y1819 (1.2 σ)
**(F)** RT: z0033 (0.8 σ), cryo: y1842 (1.2 σ)
**(G)** RT: z0089 (2 σ), cryo: y0118 (1.5 σ)
**(H)** RT: x0267 (1.5 σ), cryo: y0112 (1.5 σ)
**(I)** RT: z0115 (1.5 σ), cryo: y0660 (1.5 σ)

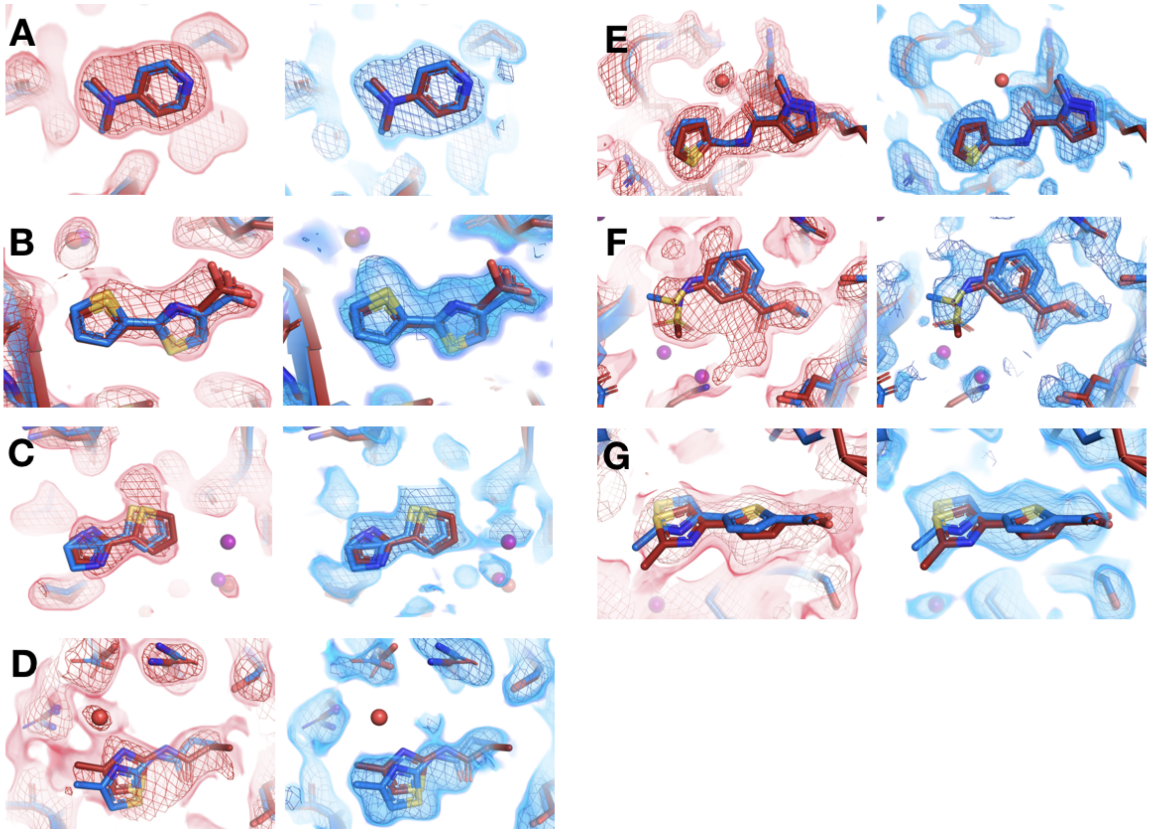


**Figure S5: All fragments that bind similarly at room vs. cryo temperatures from in-situ screen.** For each panel: RT density on left, cryo on right. Same coloring as main **Fig. 4.**

**(A)** RT: x0224 (2 σ), cryo: y0118 (1.5 σ)
**(B)** RT: x0285 (1.5 σ), cryo: y0772 (0.9 σ)
**(C)** RT: x0267 (1.5 σ), cryo: y0112 (0.9 σ)
**(D)** RT: x0228 (1.9 σ), cryo: y0076 (1.2 σ)
**(E)** RT: x0262 (1.5 σ), cryo: y1656 (1.2 σ)
**(F)** RT: x0158 (2 σ), cryo: y0175 (1 σ)
**(G)** RT: x0283 (2 σ), cryo: y0829 (1.5 σ)

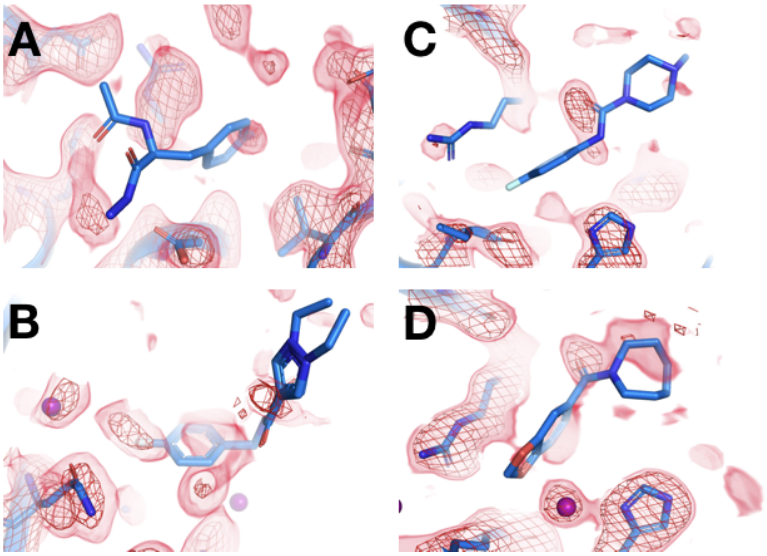


**Figure S6: Fragments that bind at cryo but not at room temperature.** Using the same 1-BDC as cryo, there is no RT density for the cryo ligand.
**(A)** RT: z0011 (1.5 σ), cryo: y0363
**(B)** RT: z0023 (2 σ), cryo: y0522
**(C)** RT: x0199 (1.8 σ), cryo: y0049
**(D)** RT: x0219 (1.5 σ), cryo: y0426

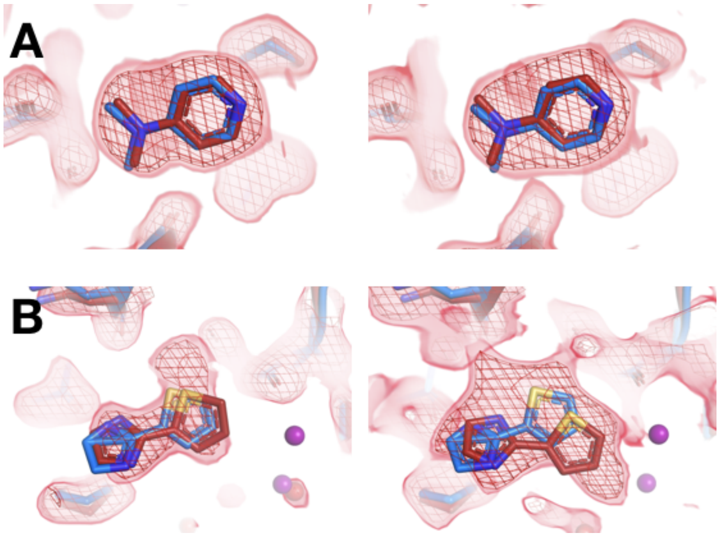


**Figure S7: Selected matching ligands across both screens have consistent binding poses.** For each panel: RT in-situ density on left, RT 1-xtal density on right. Same coloring as main **Fig. 4.**
**(A)** RT: x0224 (2 σ), cryo: y0118; RT: z0089 (2 σ), cryo: y0118
**(B)** RT: x0267 (1.5 σ), cryo: y0112; RT: z0102 (1.5 σ), cryo: y0112

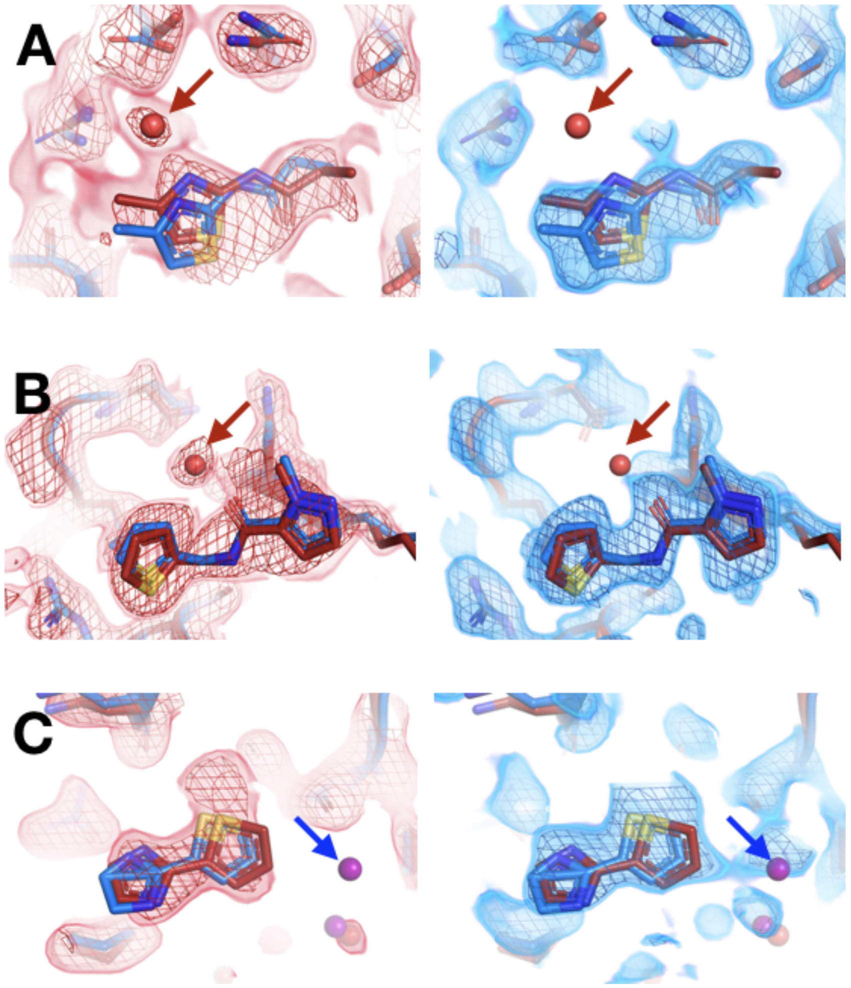


**Figure S8: Differences in solvation around fragments at room vs. cryo temperature.** Red arrows indicate a RT water with only RT event map density. Blue arrows indicate a cryo water with only cryo event map density. Typical map contour levels used when modeling are shown, but conclusions were similar when visualizing at different map contour levels. For each panel: RT density on left, cryo on right. Same coloring as main **Fig. 4.**

**(A)** RT: x0228 (1.9 σ), cryo: y0076 (1.2 σ)
**(B)** RT: x0262 (1.5 σ), cryo: y1656 (1.2 σ)
**(C)** RT: x0267 (1.5 σ), cryo: y0112 (0.9 σ)
This figure contains selected examples; see also **Fig. 6**.

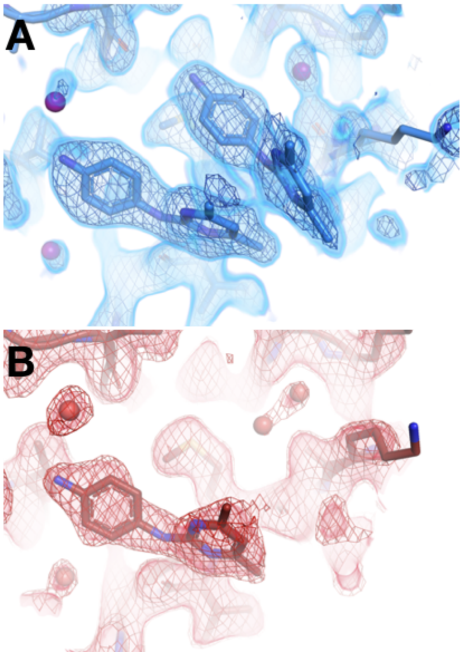


**Figure S9: Only half of a cryo stacking artifact disappears at RT.**
**(A)** Cryo y0660 (blue); cryo density (blue) 1.5 σ.
**(B)** RT z0115 (red); RT density (red) 0.7 σ.

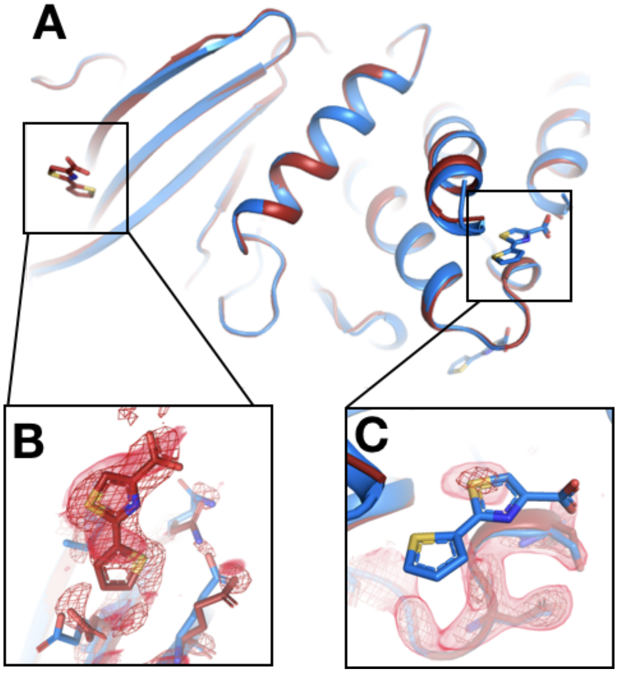


**Figure S10: Fragment that binds at a new site only at room temperature.**
**(A)** The two sites are ~40 Å away from one another.
**(B)** In the RT event map contoured at 2.5 σ (red), the RT dataset (x0285) supports a bound fragment at a new site. The cryo model (blue) from the previous cryo dataset (y0772) has no bound fragment.
**(C)** By contrast, the RT event map, calculated with 1-BDC of 0.15 (same contour), does not support the cryo model. [The site where the RT map matches the cryo site is not shown.]

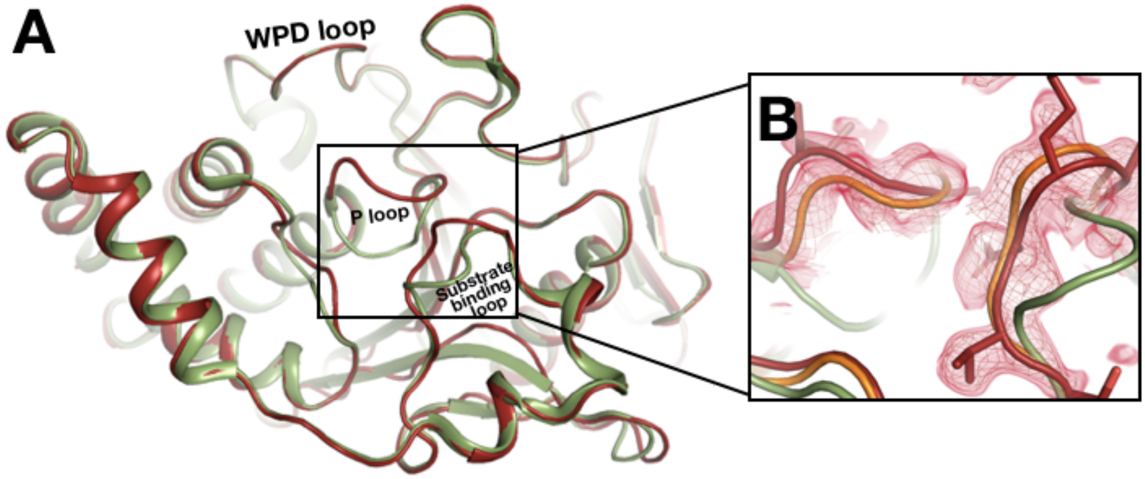


**Figure S11: Conformational difference for the active-site P loop.**
**(A)** 180° rotation around the vertical axis of main **Fig. 9A**, to view the backside of z0048 (red) aligned with 6b95 (green). The WPD loop remains open, but the P loop and the nearby substrate-binding loop adopt different conformations.
**(B)** z0048 aligned with 6b95 and a structure with Cys215 in the P loop oxidized, PDB ID 1oes (orange). RT event density is shown at 1.5 σ (red mesh). The new loop conformation at RT matches 1oes, not 6b95.

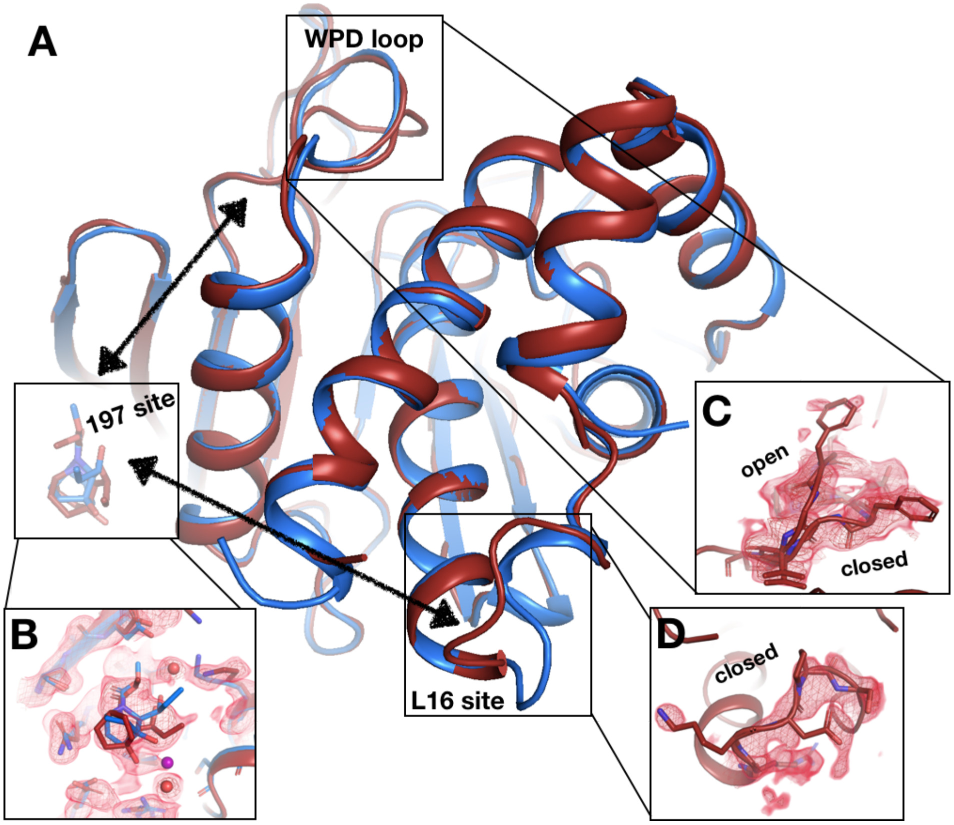


**Figure S12: Electron density evidence for allosteric protein responses at key sites seen only at RT.**
**(A)** Same as main **Fig. 11.**
**(B)** z0032 (red), y1763 (blue), RT density, 1.5 σ, indicate the fragment binds in the 197 site
**(C)** RT density, 1.5 σ, showing the WPD loop adopts alternative conformations with the WPD loop open and closed.
**(D)** RT density, 1.5 σ, is consistent with loop 16 modeled in the closed conformation.

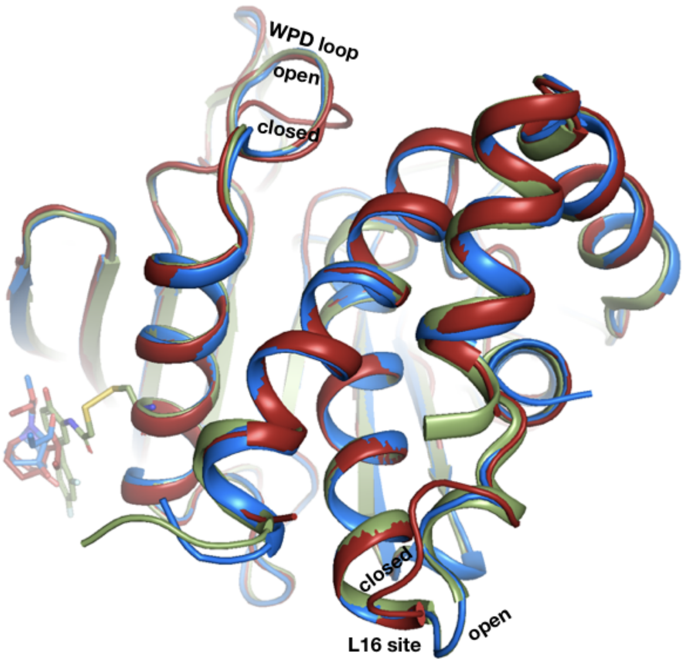


**Figure S13: Covalent allosteric inhibitor matches fragment with allosteric response at RT.** Overlay of z0032 with PDB ID 6b95, which includes a covalent inhibitor targeting the K197C mutation in the allosteric 197 site. Same coloring as in **Fig. 9**.

| | | Cluster 1 | Cluster 2 | Cluster 3 |
|---|---|---|---|---|
| Pre-merging | No. of wedges | 184 | 201 | 236 |
| | Mean resolution (Å) | 2.37 | 2.07 | 2.27 |
| | Mean a, b, c (Å) | 89.5, 89.5, 106.3 | 89.6, 89.6, 106.3 | 89.5, 89.5, 106.2 |
| | Mean $R_{merge}$ | 0.1384 | 0.1000 | 0.1242 |
| | Mean $R_{meas}$ | 0.1720 | 0.1250 | 0.1541 |
| Post-merging | No. of datasets | 41 | 40 | 47 |
| | Mean resolution (Å) | 2.17 | 1.85 | 1.98 |
| | Mean completeness | 97.2% | 99.3% | 98.0% |
| | Mean a, b, c (Å) | 89.5, 89.5, 106.3 | 89.6, 89.6, 106.3 | 89.5, 89.5, 106.2 |
| | Mean $R_{merge}$ | 0.2093 | 0.1692 | 0.2025 |
| | Mean $R_{meas}$ | 0.2289 | 0.1803 | 0.2176 |

**Table S1**: **Statistics for clusters of individual in-situ wedges, prior to merging into complete datasets.** Values shown are averages across all wedges within each cluster. See **Fig. S2** for visualization of clusters using cluster4x.

| Screen | Dataset | RT Pose Category | Notes |
|---|---|---|---|
| 1-xtal | PTP1B-z0007_x2_2 | same site, same pose | |
| | PTP1B-z0015_x2_1 | same site, same pose | |
| | PTP1B-z0021_x1_2 | same site, same pose | Found at cryo 1-BDC |
| | PTP1B-z0025_x1_1 | same site, same pose | |
| | PTP1B-z0028_x1_1 | same site, same pose | |
| | PTP1B-z0033_x2_1 | same site, same pose | Found at cryo 1-BDC |
| | PTP1B-z0089_x1_3 | same site, same pose | Also done in-situ |
| | PTP1B-z0102_x1_2 | same site, same pose | Also done in-situ |
| | PTP1B-z0115_x1_1 | same site, same pose / same site, new pose | Found at cryo 1-BDC; only 1/2 fragments bind |
| | PTP1B-z0055_x1_2 | same site, new pose | Ligand flips; active site |
| | PTP1B-z0110_x1_1 | same site, new pose* | Also done in-situ; extended Lys density |
| | PTP1B-z0032_x1_1 | same site, new pose* | Protein moves: L16 closed, WPD open and closed |
| | PTP1B-z0042_x1_1 | new site | |
| | PTP1B-z0048_x1_2 - event 3 | new site | Cryo-non-hit; extended Lys density, Lys197 |
| | PTP1B-z0048_x1_2 - event 1 | new site | Cryo-non-hit; extended Lys density, Lys237 |
| in-situ | PTP1B_is2-x0158-x0192-x0243-x0275 | same site, same pose | Found in cluster 1 (also found in all-wedges but pose was ambiguous) |
| | PTP1B_is2-x0224 | same site, same pose | |
| | PTP1B_is2-x0228 | same site, same pose | Also done in 1-xtal |
| | PTP1B_is2-x0283-x0429 | same site, same pose | Found in cluster 1 |
| | PTP1B_is2-x0285-x0431 - event 5 | same site, same pose | Found in cluster 1 |
| | PTP1B_is2-x0262-x0374 | same site, same pose | Found in cluster 2 |
| | PTP1B_is2-x0267 | same site, same pose | Also done in 1-xtal, found at cryo 1-BDC |
| | PTP1B_is2-x0260 | same site, new pose | The ligand flips |

| | | | |
|---|---|---|---|
| | PTP1B_is2-x0227-x0252 | same site, new pose | The ligand flips |
| | PTP1B_is2-x0258 | same site, new pose* | Also done in 1-xtal; extended Lys density |
| | PTP1B_is2-x0222 | same site, new pose* | Found at cryo 1-BDC; ligand binds the same but α6 helix is ordered at RT |
| | PTP1B_is2-x0256 | same site, new pose** | Ligand flips |
| | PTP1B_is2-x0285-x0431 - event 1 | new site | Found in cluster 1 |
| | PTP1B_is2-x0225 | new site | Opposite sides of the protein |

**Table S2: List of all fragment hits for room-temperature screens.** * The fragment pose is the same but the protein conformation is altered at RT vs. cryo. ** We assume the cryo pose is different but the cryo model was not published since the ligand density was poorly defined.

| | **Average MW** | **Average # rotatable bonds** | **Average # H-bonds** | **Average # non-H-bonds** | **Average iLOGP** |
|---|---|---|---|---|---|
| **new pose and/or new site (n=14)** | 196.9 | 2.36 | 1.60** | 66.0* | 1.9* |
| **same site, same pose (n=15)** | 202.5 | 2.00 | 0.73** | 43.8* | 1.5* |

**Table S3: Chemical properties of fragments and their binding sites.** H-bonds and non-H-bond interactions were calculated using Probe (Word et al., 1999); H-bonds are from ligand atoms to protein or water atoms. Other parameters were calculated using SwissADME (Daina et al., 2017). To test the significance of the difference in each parameter between the two fragment categories, Student's t-tests were performed (* $p < 0.05$, ** $p = 0.054$).

***Additional supplemental files:***

**Movie S1: Fragments in allosteric L16 site shift the α6 helix to different extents.** Fragments are shown in order of the extent to which they push the α6 helix toward the active site (up in this view) at cryo, calculated based on RMSD to the closed conformation (purple, 1sug). As the helix is pushed up, the fragments tend to bind closer to and intercalate below it, indicating a correlation between the fragment position under the helix and how much the helix is perturbed. Two RT fragments are shown, each after their corresponding cryo fragment; the effect on the helix is similar for each fragment at cryo vs. RT.
